# Comparison of methods for estimating genetic correlation between complex traits using GWAS summary statistics

**DOI:** 10.1101/2020.10.12.336867

**Authors:** Yiliang Zhang, Youshu Cheng, Wei Jiang, Yixuan Ye, Qiongshi Lu, Hongyu Zhao

## Abstract

Genetic correlation is the correlation of additive genetic effects on two phenotypes. It is an informative metric to quantify the overall genetic similarity between complex traits, which provides insights into their polygenic genetic architecture. Several methods have been proposed to estimate genetic correlations based on data collected from genome-wide association studies (GWAS). Due to the easy access of GWAS summary statistics and computational efficiency, methods only requiring GWAS summary statistics as input have become more popular than methods utilizing individual-level genotype data. Here, we present a benchmark study for different summary-statistics-based genetic correlation estimation methods through simulation and real data applications. We focus on two major technical challenges in estimating genetic correlation: marker dependency caused by linkage disequilibrium (LD) and sample overlap between different studies. To assess the performance of different methods in the presence of these two challenges, we first conducted comprehensive simulations with diverse LD patterns and sample overlaps. Then we applied these methods to real GWAS summary statistics for a wide spectrum of complex traits. Based on these experiments, we conclude that methods relying on accurate LD estimation are less robust in real data applications compared to other methods due to the imprecision of LD obtained from reference panels. Our findings offer a guidance on how to appropriately choose the method for genetic correlation estimation in post-GWAS analysis in interpretation.

## Introduction

Genetic correlation is the correlation of additive genetic effects contributing to two phenotypes. It quantifies the overall genetic similarity and provides insights into the polygenic genetic architecture of complex traits^1^. In the past 15 years, thousands of genome-wide association studies (GWAS) have been conducted, successfully identifying tens of thousands of single-nucleotide polymorphisms (SNPs) associated with complex human traits and diseases^2^. Beside association mapping for individual traits, methods have been developed to estimate genetic correlation based on linear mixed models (LMM) which use individual genotype data of independent subjects from GWAS as the design matrix^3–6^. Compared with traditional family-based approaches^7^,^8^, these GWAS-based methods do not need to collect related samples. Moreover, they do not require the studied phenotypes to be measured on the same individuals when estimating genetic correlation, which makes it possible to study a wide spectrum of human complex traits/diseases simultaneously by using different cohorts. Facilitated by the advances in genetic correlation estimation methods, genetic correlation analysis has gained popularity in the field and become a routine procedure in post-GWAS analyses. For example, a recent study performed genetic correlation analysis on 25 common brain disorders and showed that psychiatric disorders (e.g. schizophrenia and bipolar disorder) shared significantly correlated genetic risks while neurological disorders (e.g. Alzheimer’s disease and ischemic stroke) were more distinct from one another^1^. Bioinformatics servers have also been built to improve the computation and visualization of genetic correlations^9^.

Genetic correlation estimation methods can be classified as methods requiring individual-level data^6^,^10–13^ and methods that use GWAS summary statistics as input^3–5^,^14–19^. Restricted maximum likelihood (REML) is the most common approach among individual-level-data-based methods where genetic correlation is estimated as one of the (co)variance component parameters of LMM. Computational tools have been released to implement REML, such as Genome-wide Complex Trait Analysis^6,20^ (GCTA), MTG2^13^, and BOLT-REML^10^, which mainly differed by the algorithm for log-likelihood optimization. However, methods based on individual-level data have not gained as much popularity as methods based on GWAS summary statistics because of a lack of data availability and computational efficiency. Cross-trait linkage disequilibrium (LD) score regression (LDSC) is the first method that uses GWAS summary statistics alone as input to estimate genetic correlation^3^. Built upon LDSC, methods have been developed to estimate annotation-stratified^4^, local^14–16^, and trans-ethnic^17^ genetic correlation from GWAS summary statistics. Zhang et al.^15^ showed that most existing methods are based on the idea of minimizing the “distance” between the empirical and theoretical covariance matrices of marginal z-scores obtained from GWASs of two phenotypes. Due to the information loss in GWAS summary statistics, the standard errors of estimates from summary-statistics-based methods can be substantially higher than those from methods based on individual-level data^21^.

Genetic correlation analysis has a variety of downstream applications. Properly modeling genetic correlation could enhance statistical power in genetic association studies^22^,^23^, improve risk prediction accuracy^24–26^, and facilitate causal inference and mediation analysis^27–31^. However, there has been no consensus regarding which method is the most desired one to provide genetic correlation estimation under different contexts when only GWAS summary statistics are available. Although LDSC is the most widely used method in terms of genome-wide genetic correlation estimation, Lu et al.^4^ used numerical and theoretical illustrations to show that the GNOVA estimator is more accurate. A recently proposed method, high-definition likelihood (HDL)^5^, also outperformed LDSC in simulations. Although simulations were conducted to investigate the performance of these methods in each original publication, the simulation settings were largely restricted to those that could demonstrate the merits of each method. There is a need for objective and thorough analyses to benchmark and compare the performance of different genetic correlation estimation methods under realistic settings.

In this paper, we evaluate the performance of three summary-statistics-based genome-wide genetic correlation estimation methods, LDSC, GNOVA, and HDL, by comprehensive simulations and real data applications. In simulations, the phenotypes were generated using real genotype data from four different sources^32–35^. We also investigate the performance of different methods using in-sample, external, and mismatched reference panels for LD. Besides the settings satisfying the model assumptions, we evaluated the robustness of each method against model misspecification. In real data applications, we estimated genetic correlation using GWASs conducted on 10 complex traits in the UK Biobank (UKBB)^32^, 12 phenotypes in the Wellcome Trust Case Control Consortium (WTCCC) and Northern Finland Birth Cohort (NFBC), and 30 complex traits with publicly available GWAS summary statistics (**Supplementary Table 1**). Our findings provide a guidance on the statistical properties, advantages, and limitations of each method under a broad range of contexts.

## Methods

### Quality control of genotype data

In our simulations, we used imputed genotype data from four cohorts to simulate phenotypes, including UKBB (phase 3; n=487,409)^32^, WTCCC (n=15,918)^33^, NFBC (n=5,402)^34^, and Myocardial Infarction Genetics Consortium (MIGen; n=6,042)^35^. We further selected genetically unrelated 276,731 subjects of white British ancestry from UKBB in our analysis. Because genotype data in WTCCC were also gathered in Britain, WTCCC largely shares the same LD patterns with UKBB. NFBC collected genotype and health data from subjects in the two northmost provinces of Finland while subjects in MIGen were drawn from Boston, Seattle, Helsinki, Malmo, Barcelona, and Milan. Therefore, subjects in NFBC and MIGen may differ in their LD patterns with UKBB.

We restricted the analysis to autosomal variants with genotype missing rate < 0.05, imputation quality score > 0.3, Hardy-Weinberg equilibrium p-value > 1e-6, and minor allele frequency (MAF) > 0.05. We also removed all the strand-ambiguous SNPs.

### Methods for genetic correlation estimation

We considered four genetic correlation estimation methods in our comparisons, including REML^6^ which requires individual-level data, and three summary-statistics-based methods, LDSC^3^, GNOVA^4^, and HDL^5^. We included REML in the analyses to evaluate the loss of estimation precision when using summary statistics only. In the following, we briefly describe the underlying statistical framework for all these methods. Assume there are two studies with sample sizes *n*_1_ and *n*_2_, respectively, standardized trait values *ϕ*_1_ and *ϕ*_2_ follow the linear models below:

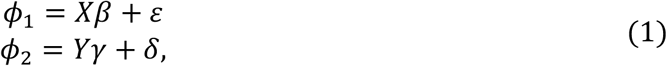

where *X* and *Y* are *n*_1_ × *m* and *n*_2_ × *m* standardized genotype matrices; *m* is the number of shared SNPs between the two studies; *ϵ* and *δ* are the noise terms; and *β* and *γ* denote the genetic effects for *ϕ*_1_ and *ϕ*_2_. The combined random vector of *β* and *γ* follows a multivariate normal distribution given by:

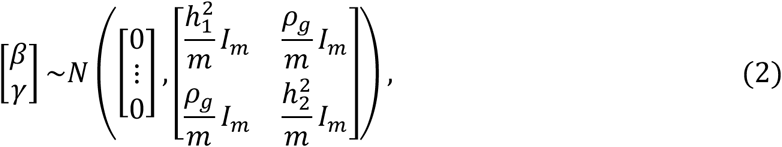

where 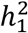 and 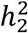 are the heritability of two traits, respectively; *ρ*_*g*_ is the genetic covariance between two traits; and *I*_*m*_ is the identity matrix of size *m*. The random effect assumption in model (2) is shared by all the methods we evaluate in this study. Without loss of generality, we assume the first *n*_*s*_ samples in each study are shared (*n*_*s*_ ≤ *n*_*1*_ and *n*_*s*_ ≤ *n*_*2*_). The non-genetic effects of the shared samples for the two traits are correlated:

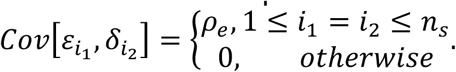

Since trait values *ϕ*_1_ and *ϕ*_2_ are standardized, we have 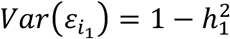 and 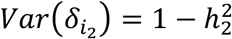 for 1 ≤ *i*_1_ ≤ *n*_1_ and 1 ≤ *i*_2_ ≤ *n*_2_. Then, genetic correlation is defined as 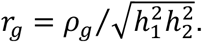

When individual-level genotype and phenotype are available, heritability and genetic covariance are estimated as the variance components of model (1) by REML. This approach has been implemented in multiple packages^10^,^13^,^20^. When only GWAS summary statistics are available, LDSC, GNOVA, and HDL can be applied for genetic correlation estimation. LDSC regresses the product of marginal z-scores in GWAS summary statistics for the two traits on LD scores of each SNP. Instead of regression, GNOVA applies a method of moments procedure. HDL calculates the joint distribution of z-scores in two GWAS and uses maximum likelihood estimation (MLE). The software of LDSC and GNOVA allow users to specify their reference panel while the reference panel to implement HDL is restricted to one of the three reference panels, UKBB imputed HapMap3 SNPs, UKBB imputed HapMap2 SNPs, and UKBB Axiom Array SNPs, provided by the software of HDL. In our simulations, we observed that REML outperformed summary-statistics-based methods^21^. In real data application on phenotypes in UKBB, WTCCC, and NFBC, since the true genetic correlations were unknown, we compared the similarity of the estimators of summary-data-based methods with REML estimator to evaluate their performance.

### Simulation settings for in-sample reference panel

In this section, the genotype data used to generate the phenotypes were also used as the LD reference panel. To implement the software of HDL, we used its provided reference panel based on 307,519 SNPs of 336,000 white British individuals from the UKBB Axiom Array data^5^. We note that both LDSC and GNOVA can use customized reference panels, hence we took the SNPs overlapping between quality controlled UKBB SNPs and UKBB Axiom Array SNPs in the HDL reference panel to construct the reference panel for LDSC and GNOVA. 305,370 SNPs (99.30% of the SNPs in HDL reference panel) and 276,731 unrelated UKBB white British individuals remained in the data. We simulated phenotypes (*ϕ*_1_ and *ϕ*_2_) based on the SNPs and samples in the reference panel. In this simulation setting, *ϕ*_1_ and *ϕ*_2_ were measured on the same set of samples (i.e. complete sample overlap).

We generated the effect sizes of SNPs by the multivariate normal distribution as in model (2) and applied GCTA^20^ to simulate *ϕ*_1_ and *ϕ*_2_. PLINK^36^ was then used to perform GWAS and obtain summary statistics of simulated traits. Following the simulation settings in Ning et al.5, we set the heritability of trait 1 and trait 2 as 0.2 and 0.4 and fixed both genetic and phenotypic correlations to be 0.5. Under this setting, all SNPs were considered causal and had equal contributions to heritability and genetic correlation. We also considered scenarios where the true correlation between traits was low or the causal SNPs of traits were sparse. We repeated each simulation setting 100 times. Detailed simulation settings are summarized below.

1. The true genetic and phenotypic correlations were set to be 0.5.
2. The true genetic correlation was set to be 0.1. The covariance of non-genetic effects (*ρ*_*e*_) was set to be 0.2.
3. The true genetic correlation was set to be 0. The covariance of non-genetic effects (*ρ*_*e*_) was set to be 0.2.
4. 30,537 out of 305,370 SNPs (10%) were randomly selected as causal variants. The true genetic and phenotypic correlations were set to be 0.5.

LDSC, GNOVA, and HDL were applied to the GWAS summary statistics. Using the same simulated individual-level GWAS data, we also conducted REML by applying BOLT-REML^10^ and compared its performance with the summary-statistics-based methods. Besides simulations on quantitative traits, we also repeated settings 1 and 3 on binary traits using a liability model. The procedure to simulate continuous liability *ψ*_1_ and *ψ*_2_ was same as the procedure to simulate *ϕ*_1_ and *ϕ*^2^. The observed trait is 1[*ψ*_*i*_ > *𝜏*], where *i* = 1,2 and *𝜏* is the 80% quantile of standard normal distribution and hence the prevalence of the binary traits were set to be 0.2.

### Simulation settings for external reference panel with matched ancestry

As a more common case in post-GWAS analysis, GWAS genotype data and reference panel are likely derived from different cohorts with a similar ancestry background (external reference panel). Here, we used the overlapping SNPs between the imputed WTCCC genotype data and the HapMap2 reference panel provided by HDL to simulate phenotypes. We chose the HapMap2 panel of HDL because it had the largest proportion of overlapping SNPs with the imputed WTCCC genotype data (85.43%; 657,181 overlapping SNPs out of 769,306 SNPs in the HapMap2 reference panel of HDL) among the three reference panels available for HDL. The HapMap2 reference panel was also used as the LD reference panel for HDL. The SNPs of the 276,731 UKBB samples in the previous section were then included in the reference panel for LDSC and GNOVA in this section.

Samples in WTCCC were randomly divided into two subgroups each with 7,959 individuals. We denote these two subgroups as set 1 and set 2, respectively. To assess how the performance of the competing methods is affected by sample overlap between GWASs, we conducted two sets of simulations for GWASs with complete sample overlaps and no sample overlap. We fixed the heritability of *ϕ*_1_ and *ϕ*_2_ as 0.5 and the value of genetic covariance was set to be 0, 0.1, and 0.2, respectively. The effect sizes of the SNPs were generated according to model (2). We also conducted simulations on spare causal SNPs where only 10% of the SNPs were randomly set to be causal SNPs. We repeated each simulation setting 100 times. Details on simulation settings are summarized below.

1. Complete sample overlap: *ϕ*_1_ and *ϕ*_2_ were both simulated on set 1. The covariance of non-genetic effects on shared samples was set to be 0.2.
2. No sample overlap: we used set 1 and set 2 to simulate *ϕ*_1_ and *ϕ*_2_, respectively.

We also simulated binary traits using the liability model described in the previous section and compared the performance of different methods on binary traits. Next, following the same procedure, we performed additional simulations for HDL using the SNPs before quality control in imputed WTCCC genotype data. We only excluded ambiguous or multiallelic SNPs. There were 769,236 overlapping SNPs out of the 769,306 SNPs (99.99%) in the HapMap2 reference panel of HDL.

### Simulation settings for external reference panel with mismatched LD

We investigated the robustness of LDSC, GNOVA and HDL in the situation that the GWAS samples are from a population distinct from the reference panel population, i.e., the LD patterns differ between the GWAS population and the reference panel population. Here, we used the imputed genotype data from NFBC and MIGen to simulate *ϕ*_1_ and *ϕ*_2_, respectively. We still used the HapMap2 reference panel for HDL. After qualify control, there were 745,288 (96.88% of the SNPs in the HapMap2 reference panel of HDL) overlapping SNPs in the imputed NFBC and MIGEN genotype data and SNPs in HapMap2 reference panel, which were taken forward to generate *ϕ*_1_ and *ϕ*_2_. Similarly, the UKBB genotype data were still used as the LD reference panel for LDSC and GNOVA.

Based on model (2), the heritability for both phenotypes was set to be 0.5 and the genetic covariance was set to be 0, 0.1, and 0.2, respectively. We first assumed infinitesimal model where all the SNPs were causal SNPs. Then, we generated the phenotypes based on 10% randomly selected SNPs from all the SNPs. We compared the performance of the competing methods on the GWASs summary data calculated from the simulated phenotypes. Each simulation setting was repeated for 100 times. We also compared the performance of the three methods on binary traits.

### Genetic correlation estimation for phenotypes in UKBB, WTCCC and NFBC

We compared the performance of LDSC, GNOVA, and HDL on the phenotypes in UKBB, WTCCC, and NFBC where individual-level data are available, so that we could also compare the differences between summary-statistics-based estimators and the more accurate REML estimator^21^. The UKBB genotype data were still used as the reference panel for the summary-statistics-based methods. Sex, age, and top 4 principal components were included as covariates to perform GWAS. We included both quantitative and binary traits in our analyses. All methods we applied here provided justification for applications on binary traits^3–6^.

We first applied the methods to estimate genetic correlations across 10 common phenotypes in the UKBB where we extracted the genotype and phenotype data for white British individuals. Note that we are using in-sample reference panel for the UKBB phenotypes. We used the same set of SNPs included in our simulation for in-sample reference panel to perform GWAS, which accounted for 99.30% of the SNPs in the genotype array reference panel of HDL. The details of the phenotypes and the sample sizes from the UKBB dataset are summarized in **Supplementary Table 2**.

Then we further applied the methods to estimate the genetic correlations across 12 phenotypes in WTCCC and NFBC datasets. Note that in this case we used the UKBB genotype data as the external reference panel, which corresponds to matched LD structure for phenotypes from WTCCC and mismatched LD structure for phenotypes from NFBC. We note that there was no shared sample between WTCCC and NFBC. We used the same set of SNPs included in our simulation for WTCCC and NFBC to perform GWAS, which accounted for 99.99% and 96.88% of the SNPs in the HapMap2 reference panel of HDL, respectively. The details of the phenotypes and sample sizes are summarized in **Supplementary Table 3-4**.

### GWAS summary statistics of 30 complex traits

GWAS summary statistics of 29 complex traits included in our analyses are publicly available. We obtained the summary statistics of a recent lung cancer GWAS directly from the authors^37^. All GWASs were conducted on samples of primarily European descent. Our choice of these traits was based on the preference for larger sample sizes and higher SNP overlapping rates with the HDL reference panel. The HapMap3 reference panel of HDL has 1,029,876 SNPs. More than 90% of the SNPs in the HDL reference panel also showed up in the GWASs of 26 of the 30 complex traits. Suicide attempt (SA) had the least overlapping SNPs with the HDL reference panel (81.81%). For fairness, the reference panel for LDSC and GNOVA was also constructed by the 1,160,014 HapMap3 SNPs in our UKBB dataset. The details about the sample sizes and the sources of the 30 traits are given in **Supplementary Table 1**.

## Results

### Simulation results

We compared the performance of LDSC, GNOVA, and HDL on point estimation of genetic covariance and correlation, type I error control, and statistical power. To investigate the robustness of these methods to the choice of LD reference panel, we performed simulations on in-sample reference panel, external reference panel with matched LD, and external reference panel with mismatched LD. Unlike LDSC or GNOVA, which can use customized reference panel, the software of HDL provides the pre-calculated eigenvalues and eigenvectors of LD matrix and restricts the reference samples to the UKBB samples. So, for fairness, the reference panel for LDSC and GNOVA was also based on the white British individuals in UKBB. We repeated each setting 100 times.

We first followed the simulation settings in Ning et al.^5^ and simulated phenotypes based on the genotype data in the reference panel (i.e. in-sample reference panel) constructed by the UKBB samples. A total of 99.30% of the SNPs in the HDL reference panel were used to simulate the phenotypes. We also compared the difference between the summary-statistics-based methods and REML^6^. As expected, REML outperformed all summary-statistics-based methods even when these methods were provided with the in-sample reference panel. Both LDSC and GNOVA provided unbiased estimates for genetic covariance (Figure 1A) in all settings. LDSC achieved more accurate estimation than GNOVA. By contrast, HDL overestimated genetic covariance when the true genetic covariance was relatively high. Nevertheless, estimates of HDL had the least variance among the summary-statistics-based methods and were more accurate under weak genetic covariance. For genetic correlation estimation, all methods including HDL provided unbiased estimates (Figure 1B). Due to the ratio form of genetic correlation, the bias of HDL in genetic covariance and heritability estimation was cancelled out. Compared with other summary-statistics-based methods, HDL showed the best performance on genetic correlation estimation and showed the least difference with REML. Though achieving comparable power with REML, HDL had the largest type I error. GNOVA showed the largest variance for genetic correlation estimation and the lowest statistical power and type I error (**Supplementary Figure 1**). All methods were robust to model misspecification where 10% SNPs were set as the causal SNPs (**Supplementary Figure 2**).

**Figure 1.**
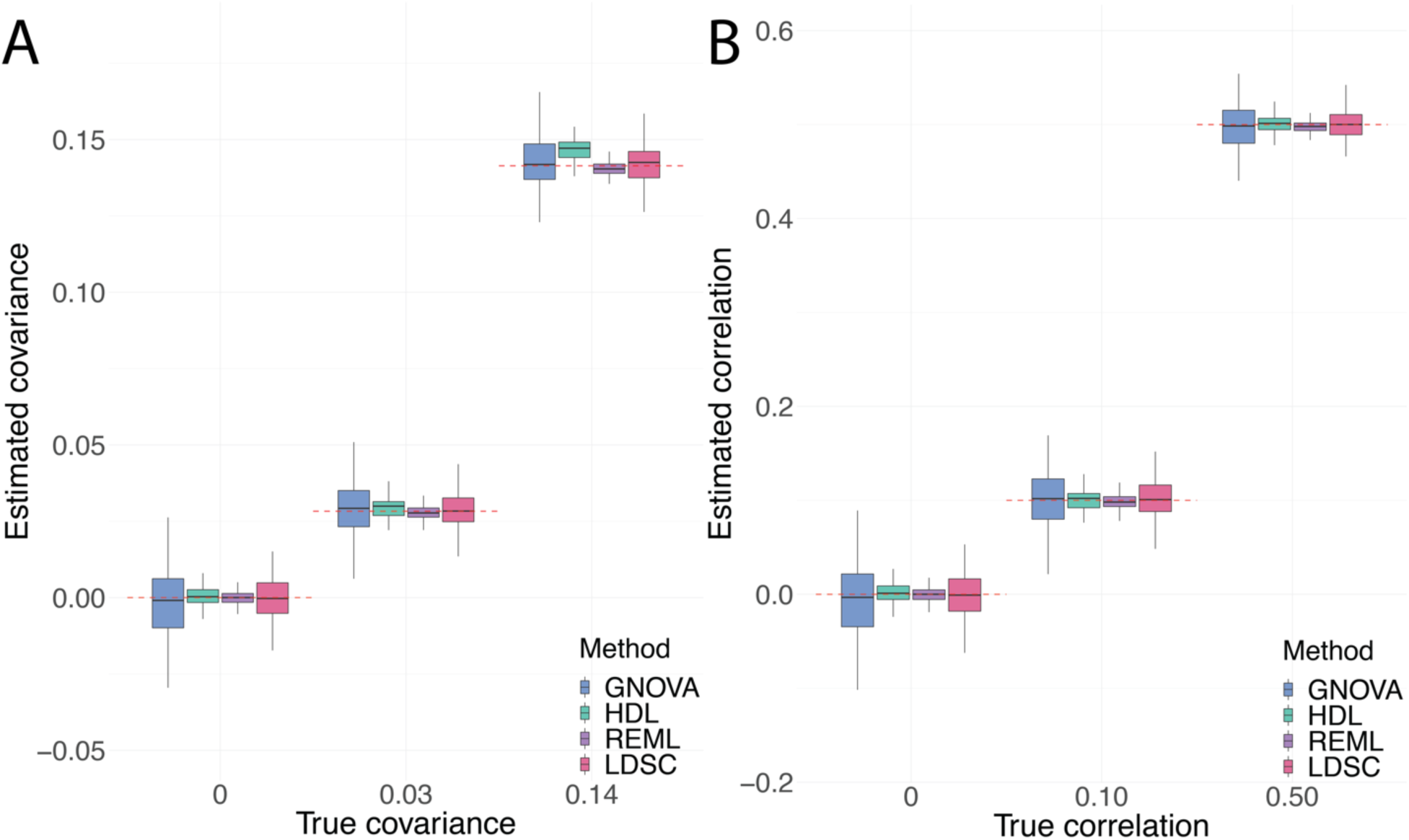
Comparisons of genetic covariance and correlation estimation using in-sample reference panel. The estimates for **(A)** genetic covariance and **(B)** genetic correlation among LDSC, GNOVA, HDL and REML are demonstrated by boxplot which shows the quantiles of the estimates. The red dashed lines represent true values.

Next, we used the genotype data in WTCCC to generate phenotypes. The genotype data in WTCCC were collected across the UK and should share a similar LD structure with UKBB (external reference panel with matched LD). There were 85.43% of the SNPs in HDL reference panel also appeared in the imputed WTCCC genotype data after quality control, which were used to generate the phenotypes. When the two GWASs were simulated on the same group of samples (set 1), LDSC and GNOVA could provide similar unbiased estimates for both genetic covariance and correlation. HDL consistently overestimated the parameters, except for estimating genetic correlation when the true value was relatively large (Figure 2A-B), where the bias in the estimation of heritability and genetic covariance was cancelled out. There was severe type I error inflation for HDL while GNOVA achieved the lowest type I error across the competing methods and comparable power with LDSC (**Supplementary Figure 3**). The bias and type I error inflation of HDL in GWAS with complete sample overlap suggested that HDL could not distinguish genetic covariance from technical covariance. On the other hand, when there was no overlapping sample between the two studies (set 1 and set 2), LDSC and GNOVA still presented consistent estimates while HDL showed biased estimates, although the variance of HDL estimator was consistently lower than the estimators of LDSC and GNOVA (**Supplementary Figure 4**). No method showed inflated type I error rate and LDSC showed the least type I error. The three methods showed similar statistical power (**Supplementary Figure 5**). The results were similar under model misspecification (**Supplementary Figure 6**). We performed additional simulations to evaluate HDL by using the imputed genotype before quality control from WTCCC, where we only excluded ambiguous or multiallelic SNPs and kept the SNPs overlapping with the HDL reference panel. Although 99.99% of the SNPs in the HDL reference panel were included for the additional simulation, HDL still could not adjust for sample overlap (**Supplementary Figure 7**). However, in the additional simulation, we observed that HDL provided more accurate estimates for the parameters on GWAS without sample overlap compared with its performance using the WTCCC SNPs after quality control (85.43% overlapping SNPs) (**Supplementary Figure 4**). This indicated that the performance of HDL can be affected by the choice of SNP set in GWAS (**Supplementary Figure 7**).

**Figure 2.**
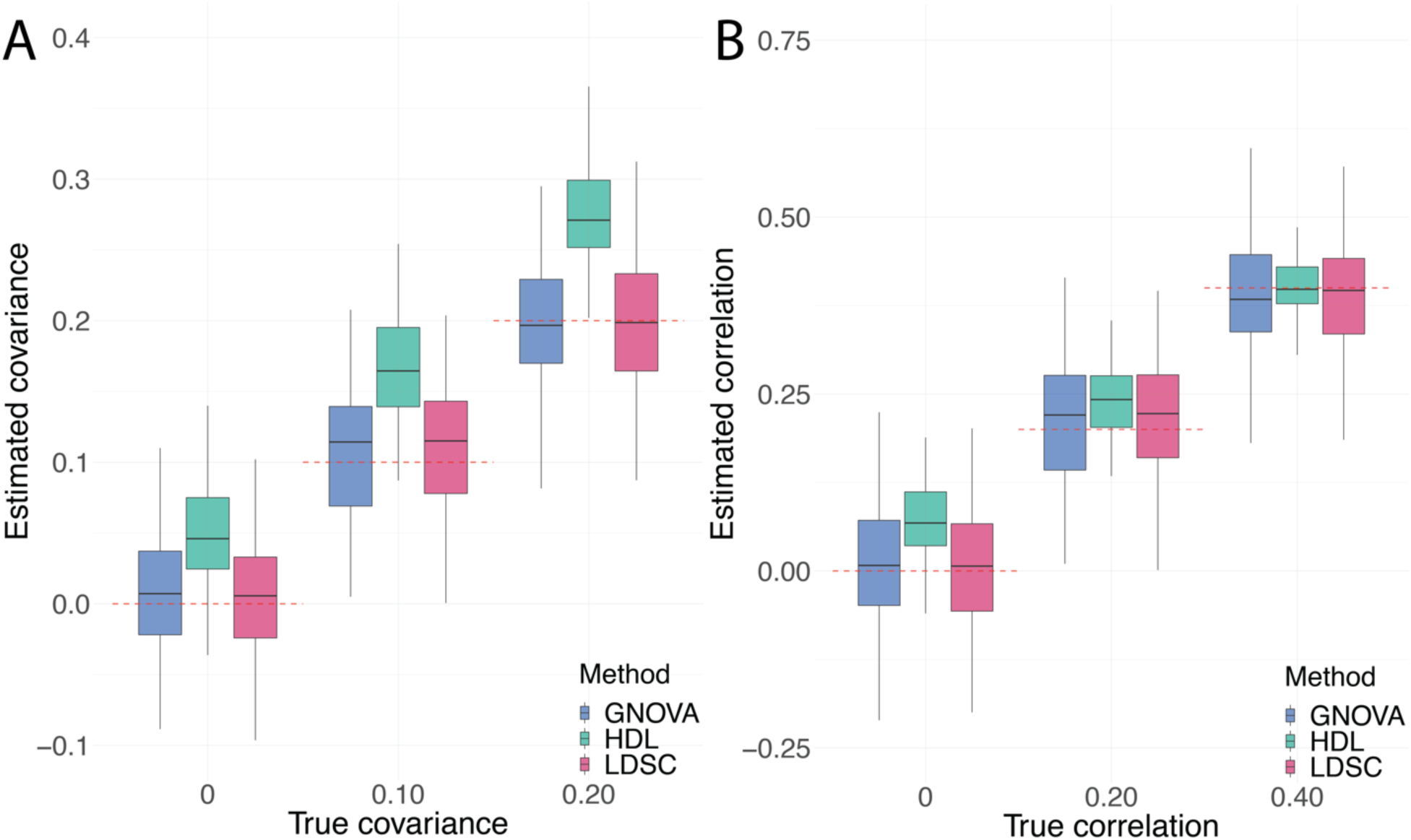
Comparisons of genetic covariance and correlation estimation using external reference panel with matched LD. We compare the estimation of genetic **(A)** covariance and **(B)** correlation when the two GWASs were simulated on the same dataset with a 100% sample overlap. The red dashed lines represent true values.

Finally, we examined the robustness of LDSC, GNOVA and HDL to the reference panel with mismatched LD. In practice, if the GWAS is conducted on populations without an ideal LD reference panel, researchers might have to use mismatched reference panel for post-GWAS analysis. We simulated *ϕ*_1_ for 5,402 samples from northern Finland in NFBC and *ϕ*_2_ for 6,042 samples across north America and Europe in MIGen. There were 96.88% SNPs in the HDL reference panel used to simulate the phenotypes. All methods showed unbiased estimators for genetic covariance except the estimation of HDL for high genetic covariance. Both LDSC and GNOVA underestimated the genetic correlation while HDL, on the contrary, overestimated the genetic correlation (Figure 3).

**Figure 3.**
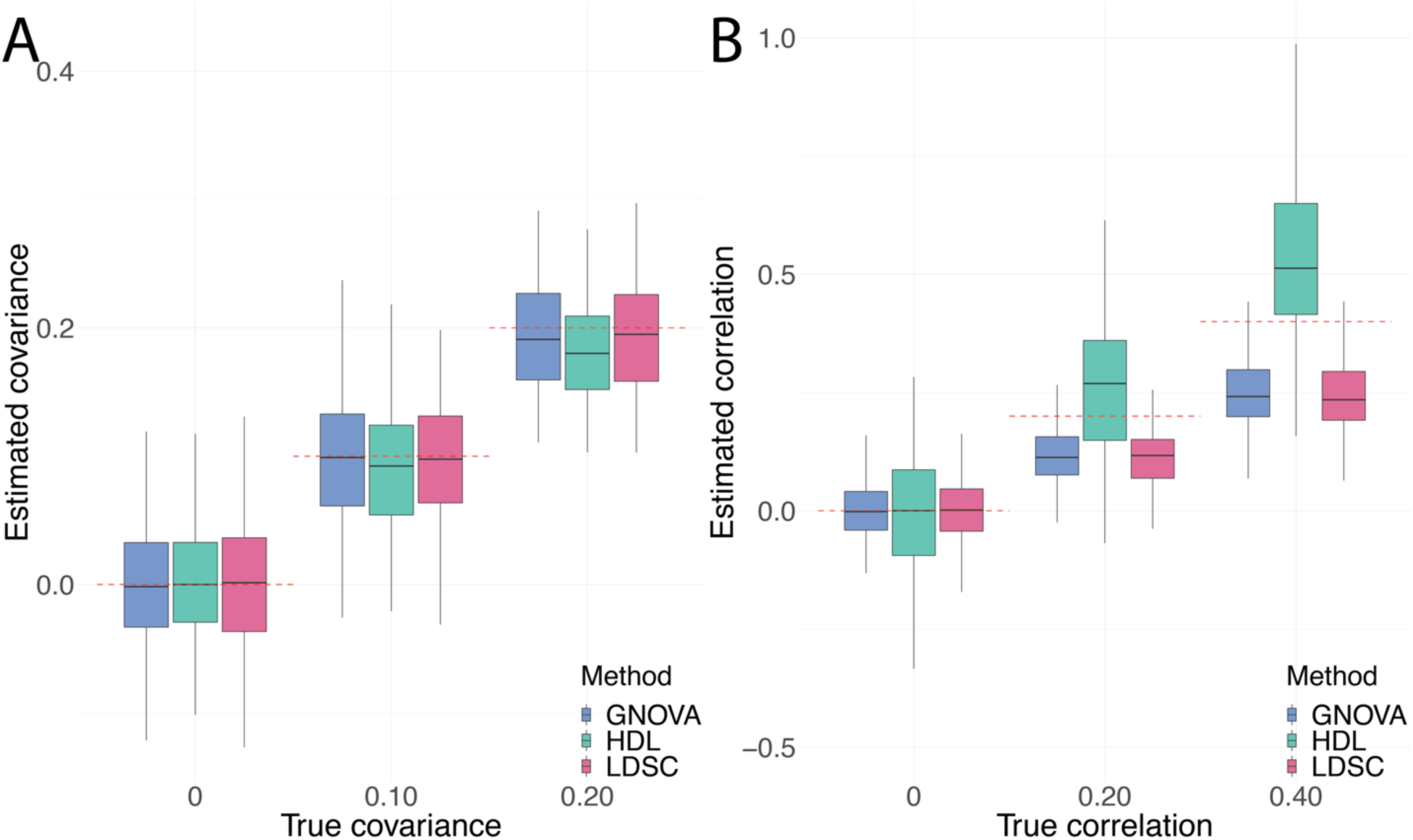
Comparisons of genetic covariance and correlation estimation using external reference panel with mismatched LD. The estimates for **(A)** genetic covariance and **(B)** genetic correlation among LDSC, GNOVA and HDL are demonstrated by boxplot which shows the quantiles of the estimates. The red dashed lines represent true values.

So, genetic covariance estimation was more stable than genetic correlation estimation under mismatched LD across the three methods. The results were similar under model misspecification (**Supplementary Figure 8**). All three methods showed moderate type I error inflation and similar power (**Supplementary Figure 9**). Compromised by false discovery issues, the results of summary-statistics-based methods should be cautiously interpreted when applied to mismatched reference panel with GWAS.

All these methods can be applied to binary traits. Although the genetic covariance is estimated on the observed scale^3^, the estimates for genetic correlation of binary traits were consistent to the estimates for quantitative traits except for larger variations across the 100 repeats in each setting (**Supplementary Figures 10-12**). There was a loss of statistical power of the methods from quantitative traits to binary traits due to less effective sample sizes (**Supplementary Figures 13-16**).

### Application to summary statistics from UKBB, WTCCC and NFBC

We compared the performance of competing methods on GWAS with individual-level data so that we could also implement and compare with REML. We used traits in UKBB, WTCCC, and NFBC which can be seen as the counterparts for in-sample reference panel, external reference panel with matched LD, and external reference panel with mismatched LD in our simulation. We included both quantitative and binary traits in our analyses. For binary traits, genetic covariance estimation is on the observed scale but there is no distinction between observed- and liability-scale genetic correlation3. In addition, we have shown through simulations that REML also outperformed summary-statistics-based methods in genetic correlation estimation for binary traits. Given these considerations, since we do not know the true genetic correlation in real data, the performance of different summary-statistics-based methods was assessed by the difference from REML results.

We first estimated genetic correlation among 10 common traits from UKBB (**Supplementary Table 2**). After Bonferroni correlation (p < 0.05/45 = 1.11e-3), 31 out of 45 pairs were identified by REML, while LDSC, GNOVA, and HDL identified 25, 22 and 22 trait pairs, respectively (**Supplementary Table 5**). All the trait pairs identified as significantly correlated by summary-statistics-based methods were also identified by REML (Figure 4A). The estimators of LDSC, GNOVA, and HDL showed considerable similarity with REML estimators with R square 0.99, 0.98, 0.99, respectively (**Supplementary Figure 17**). We note that the absolute values of most of the genetic correlations among the 45 trait pairs were less than 0.5, similar to the settings where HDL outperformed LDSC and GNOVA in our simulations.

**Figure 4.**
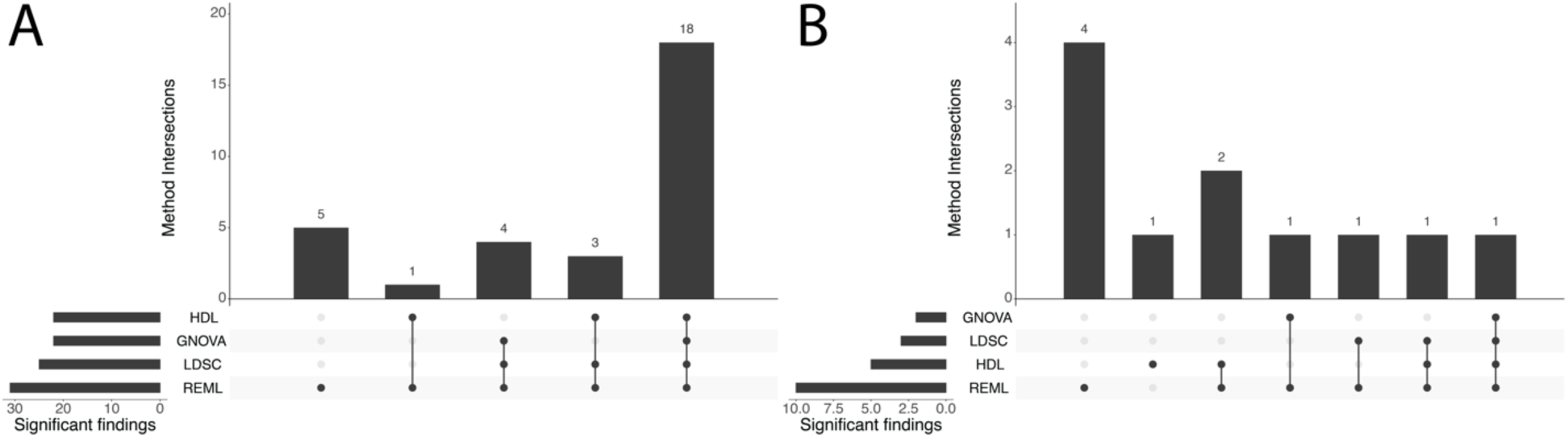
Trait pairs with significant genetic correlation identified by LDSC, GNOVA, HDL and REML for real GWAS data in UKBB, WTCCC, and NFBC. This plot uses bars to break down the Venn diagram of overlapped regions in different categories. The four categories shown in the lower panel are correlated trait pairs in **(A)** UKBB (in-sample reference panel) and **(B)** WTCCC + NFBC (external reference panel) identified by LDSC, GNOVA, HDL, and REML.

We then estimated genetic correlation across 12 traits in WTCCC and NFBC using the external reference panel. After Bonferroni correlation (p < 0.05/66 = 7.58e-4), 10 out of 66 pairs were identified by REML, while LDSC, GNOVA, and HDL identified 3, 2 and 5 trait pairs, respectively (**Supplementary Table 6**). GNOVA was the most conservative method in this analysis. We observed that all the trait pairs reported by LDSC or GNOVA were also reported by REML. However, the genetic correlation of one trait pair, hypertension (HT) and Crohn’s disease, which was not identified by REML (p=0.24), was identified by HDL (p = 3.4e-4; Figure 4B). There was a caveat of false discovery for this trait pair, because HDL failed to adjust for non-genetic covariance and suffered from severe type I error inflation with external reference panel and complete sample overlap in our simulation. Due to limited sample size and information loss because of using external reference panel, less consistency with REML results was shown by the summary-statistics-based estimators (**Supplementary Figure 18**). LDSC showed the highest correlation with REML in genetic correlation estimation compared with other methods (R square = 0.66).

### Genetic correlation of 30 complex traits

We applied LDSC, GNOVA, and HDL to estimate genetic correlation among 30 complex traits including neuropsychiatric disorders, immune diseases, cardiovascular diseases, cancer, anthropometric traits, and metabolic traits for European population. We summarized detailed information about each trait, including abbreviations and data sources, in **Supplementary Table 1**. There were 126, 112, and 166 trait pairs identified by LDSC, GNOVA, and HDL, respectively (p < 0.05/435 = 1.15e-4; **Supplementary Figure 19** and **Supplementary Table 7**). 104 trait pairs were identified by all three methods and 42 trait pairs were exclusively identified by HDL (**Supplementary Figure 20**). The point estimates for genetic correlation showed consistency across three methods (Figure 5). The estimates of LDSC and GNOVA were more similar to each other compared with HDL (R square = 0.91). Three trait pairs, including body mass index (BMI) and Hemoglobin A1(C) (HBA1C), coronary artery disease (CAD) and HBA1C, and type-II diabetes (T2D) and HBA1C, showed high genetic correlation estimation across the three methods were identified as significantly correlated by LDSC and GNOVA but were not identified by HDL (Figure 5B-C and **Supplementary Figure 21**). HBA1C measures the long-term blood glucose concentrations and has been widely used as a diagnostic test for T2D^38^,^39^. The positive associations between HBA1C and CAD has long been observed in epidemiological studies^40^. A recent Mendelian Randomization analysis further confirmed the causal role of HBA1C in developing CAD^41^. Obesity (BMI) is also known to be correlated with poor control of HBA1C and T2D^42^.

**Figure 5.**
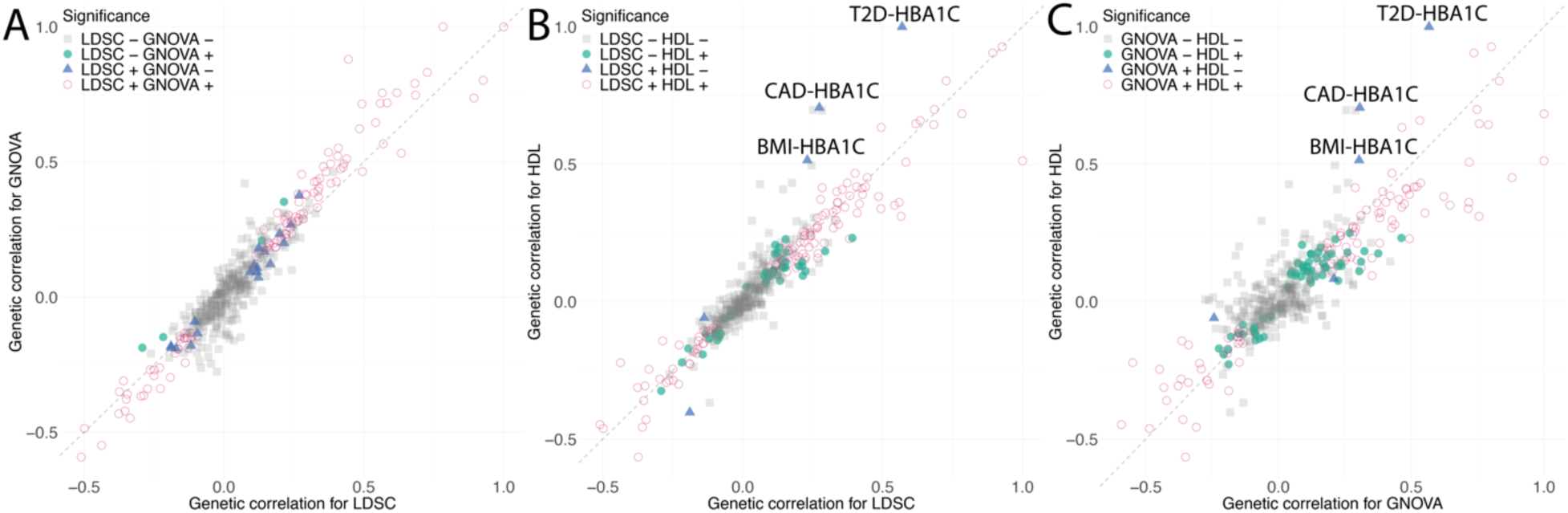
Comparisons of point estimates of genetic correlation among LDSC, GNOVA, and HDL. The comparisons are presented by scatter plots for **(A)** LDSC vs. GNOVA, **(B)** LDSC vs. HDL, and **(C)** GNOVA vs. HDL with R square 0.91, 0.85, 0.73. Each point represents a trait pair. Color and shape of each data point denote the significance level. The grey dashed lines are *y* = *x*.

## Discussion

Owing to increasingly accessible GWAS summary statistics and advances in statistical methods, genetic correlation estimation has become a routine procedure in post-GWAS analyses. The statistical challenges of summary-statistics-based methods are mainly reflected in two technical issues: (1) extensive LD between SNPs; and (2) pervasive sample overlap across GWASs. The utility of a method is largely determined by the ability to adjust for LD and overlapping samples. Through a plethora of simulations and real data applications, we have demonstrated that LDSC and GNOVA were more similar methods and robust to LD and sample overlap compared with HDL.

Because GWAS summary statistics are constructed by marginal regression statistics, LD between SNPs is embraced in the covariance of marginal effect sizes which impede us from directly estimating genetic correlation by calculating the correlation between marginal effect sizes. When only GWAS summary statistics are available, we still need additional individual-level genotype data as the reference panel to adjust for LD even for summary-statistics-based methods. So, the choice of reference panel is central to their performances. We used different types of reference panel, including in-sample reference panel and external reference panel with or without matched LD to investigate the influence of reference panel. For in-sample reference panel, GWAS and reference samples are from the same samples where the information loss of summary-statistics-based methods is minimal. However, we demonstrated that REML was actually a better choice for both quantitative and binary traits when individual-level phenotype and genotype data in GWAS are available if computational efficiency is not a concern. For external reference panel with matched LD, GWAS and reference samples are independent but share the same LD structure, i.e. are from the same population. For example, UKBB genotype data can be used as matched reference panel for GWAS performed on white British individuals. In practice, this is the most common case that the access to individual information from GWAS dataset is limited due to logistical challenges in data sharing. For reference panel with mismatched LD, GWAS and reference samples are from different populations. This can happen when the GWASs are conducted on minority population which may not have corresponding reference panel. To estimate genetic correlation, researchers have to use the reference panel from another population with similar LD structure which might lead to bias in the estimation of LD scores.

Two GWASs often share a subset of samples especially for meta-analyses. For example, part of the control samples of Grove et al.^43^ for autism spectrum disorder (ASD) and Demontis et al.^44^ for attention deficit/hyperactivity disorder (ADHD) were collected on the same cohort from Denmark by the Lundbeck Foundation Initiative for Integrative Psychiatric Research45 (iPSYCH). The exact number of overlapping sample sizes is often unavailable for large GWAS meta-analyses. We compared the performance of LDSC, GNOVA, and HDL on GWAS summary data with or without sample overlap to assess their ability to distinguish genetic correlation from non-genetic correlation.

LDSC and GNOVA can provide unbiased estimation for genetic covariance across all the simulation settings, except for in-sample reference panel where the estimates of GNOVA were sub-optimal. Overall, LDSC and GNOVA had similar performance. Both methods presented relatively consistent estimators with REML when applied to real GWAS data. On the other hand, HDL is sensitive to the choice of the reference panel because HDL used more information from the LD matrix to implement MLE^5^ but the procedure to select eigenvalues and eigenvectors of the LD matrix did not consider potential departure from the LD matrix in the GWAS data to be analyzed. Hence the results were not robust. Genetic covariance estimation of HDL was biased in most settings. The only setting in our simulation where HDL could provide unbiased estimator of genetic covariance was that 99.99% of the SNPs in the reference panel of HDL were included in the two GWAS without any sample overlap. Due to the ratio form, the genetic correlation estimation of HDL was more reliable than genetic covariance estimation and outperformed LDSC and GNOVA with in-sample reference panel. However, when applied on external reference panel, the genetic correlation estimation of HDL was unstable and had type I error inflation in simulations. For computational reasons, the software of HDL restricted the users to three reference panels but the method is not robust to the SNP set in GWAS. Therefore, in real data applications, we caution the users for potential false discovery of HDL especially for GWASs with overlapping samples. There was also significant deterioration of performance when the SNPs in GWAS decreased from 99.99% to 85.43% of the SNPs in the refence panel. In comparison, there is more flexibility of choosing the reference panel for LDSC and GNOVA by the users. None of the methods worked well for genetic correlation estimation under a mismatched reference panel. Therefore, it is always crucial to have an appropriate reference panel for summary-statistics-based methods. However, genetic covariance can be a more robust quantity to estimate with a mismatched reference panel compared with genetic correlation. Simulations under model misspecifications showed that all the methods were robust to sparsity of causal SNPs.

In summary, we have evaluated three summary-statistics-based genetic correlation estimation methods using simulations and real data applications. We compared the robustness of the methods to LD estimation and sample overlap. However, our study has several limitations. First, genome-wide genetic correlation only reflects the average concordance of genetic effects across the genome and often fails to reveal the stratified, heterogenous pleiotropic effects, especially when the underlying genetic basis involves multiple etiologic pathways^15^. Second, Speed et at.^18^ introduced a statistical framework to estimate heritability where allelic effects are function of LD and MAF. However, we restricted the methods compared in this paper established on model (2). It is possible that a more general model can also improve genetic correlation estimation. Third, there is no gold standard to compare the methods in real data application as the true genetic correlation between trait pairs is unknown. Downstream analyses of genetic correlation estimation such as multi-trait association mapping^22^, genomic structural equation modeling^23^ (GenomicSEM) and Mendelian randomization^27^,^28^ might help assess the performance of these methods in real data applications.

