## Supplementary Figures for "Comparison of methods for estimating genetic correlation between complex traits using GWAS summary statistics"

**Supplementary Figure 1. Evaluation of type I error control and statistical power among genetic correlation (covariance) estimation methods using in-sample reference panel. (A)** Type I error and statistical power when true parameters are zero or nonzero, respectively. **(B)** Type I error when the true genetic correlation is 0. **(C)** qq-plot of p value when the true genetic correlation is set to be 0.

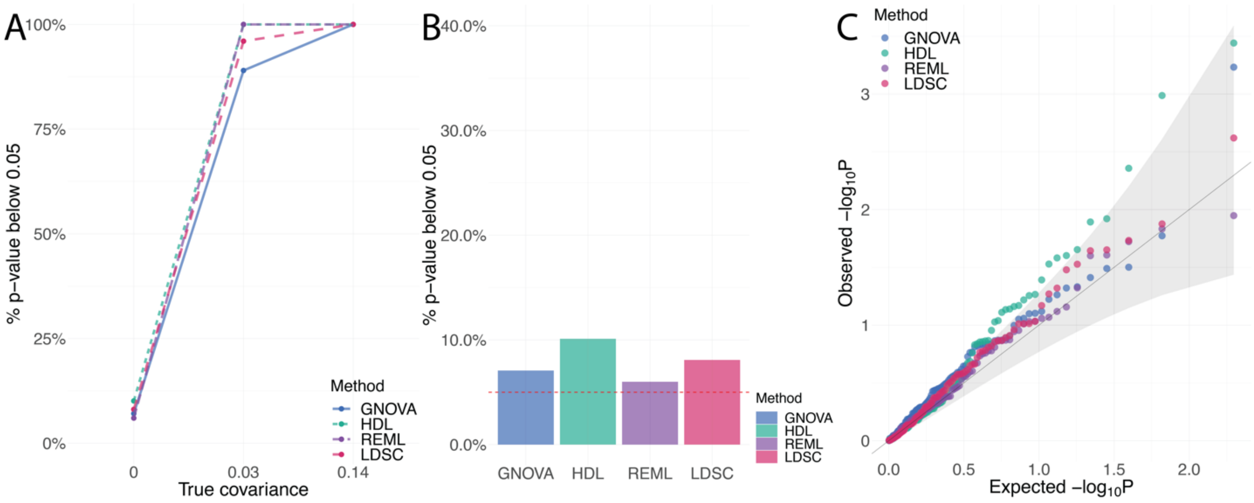

**Supplementary Figure 2. Comparisons of genetic covariance and correlation estimation using in-sample reference panel under model misspecification.** We set 10% of the total SNPs as the causal SNPs to simulate the phenotypes. The estimates for **(A)** genetic covariance and **(B)** genetic correlation among LDSC, GNOVA, HDL, and REML are summarized by boxplots which show the quantiles of the estimates. The red dashed lines represent true values.

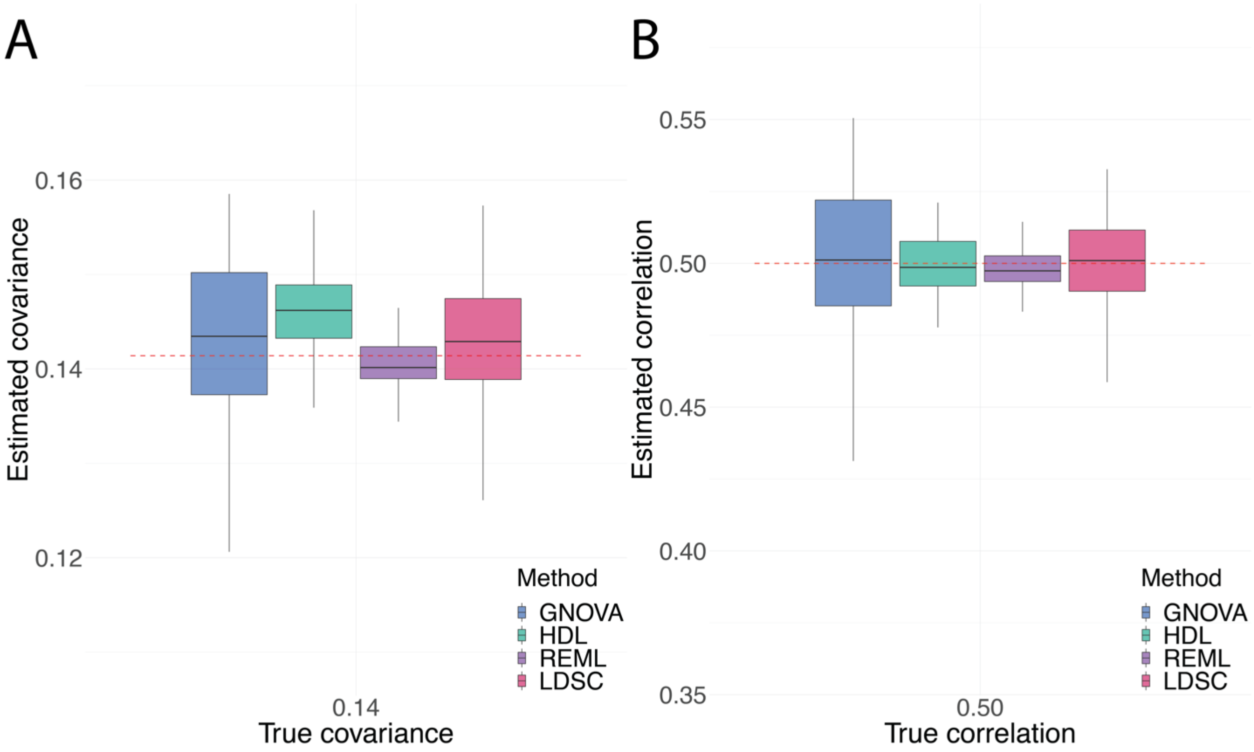

**Supplementary Figure 3. Evaluation of type I error control and statistical power among genetic correlation (covariance) estimation methods using external reference panel with matched LD on completely overlapping dataset (set 1). (A)** Type I error and statistical power when true parameters are zero or nonzero, respectively. **(B)** Type I error when the true genetic correlation is 0. **(C)** qq-plot of p value when the true genetic correlation is set to be 0.

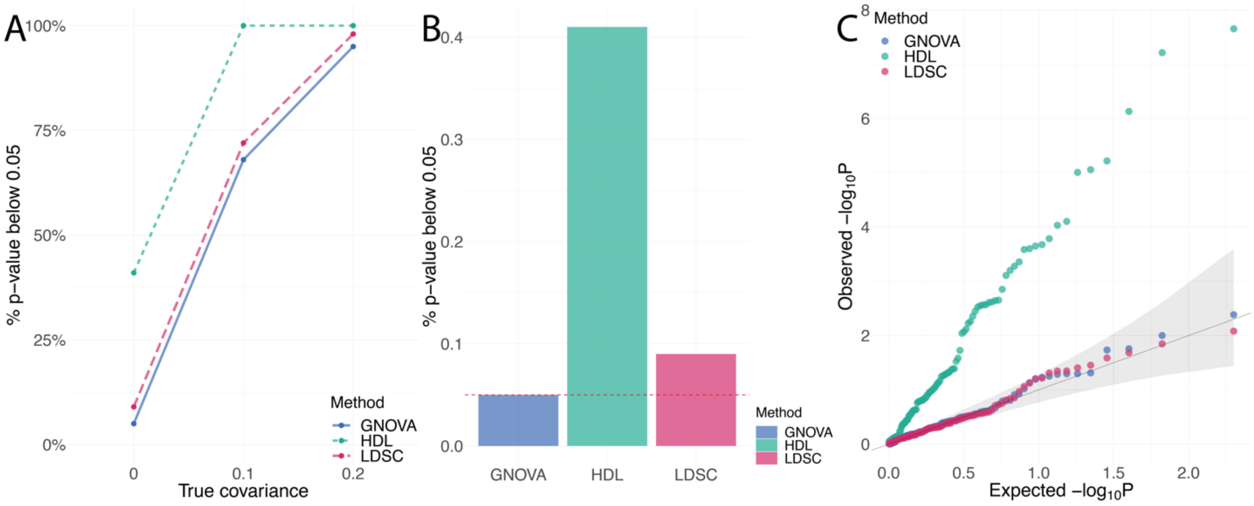

**Supplementary Figure 4. Comparisons of genetic covariance and correlation estimation using external reference panel with matched LD when the GWASs were simulated on two non-overlapping datasets.** The estimates for **(A)** genetic covariance and **(B)** genetic correlation among LDSC, GNOVA and HDL are summarized by boxplots which show the quantiles of the estimates. We compare the methods when the GWASs were simulated on two non-overlapping datasets. The red dashed lines represent true values.

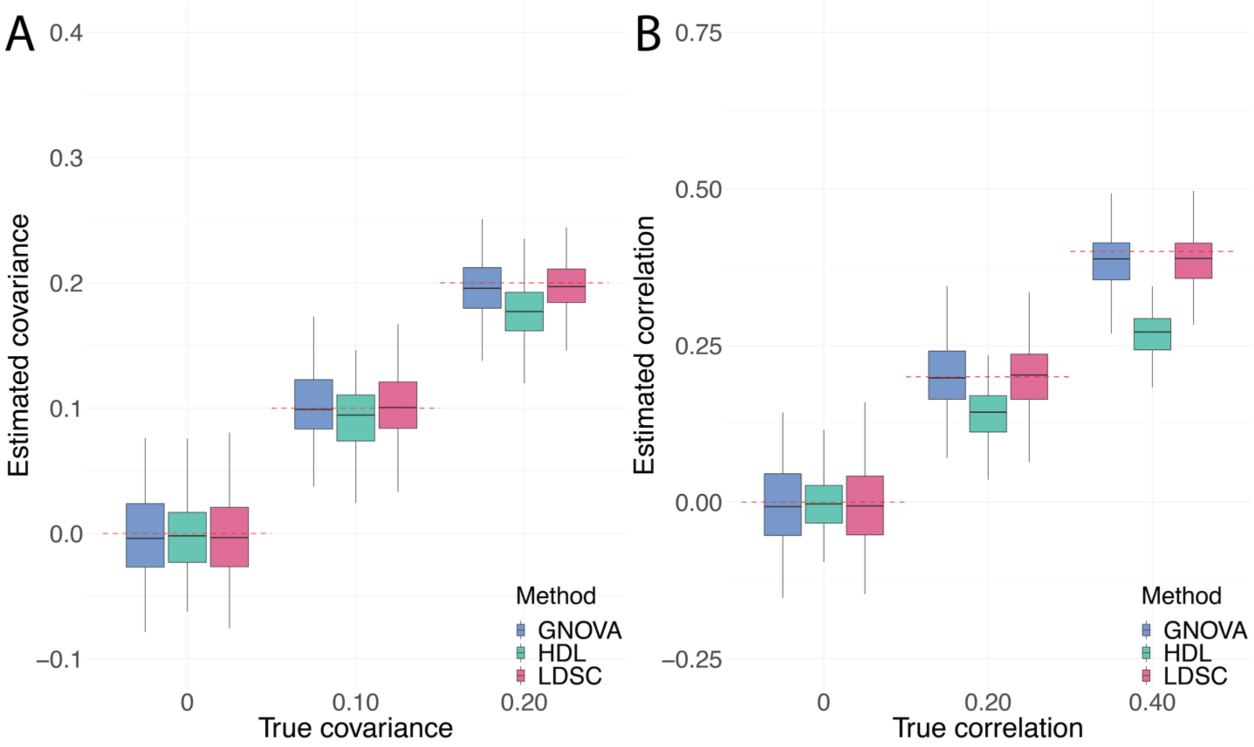

**Supplementary Figure 5. Evaluation of type I error control and statistical power among genetic correlation (covariance) estimation methods using external reference panel with matched LD on non-overlapping dataset (set 1 and set 2). (A)** Type I error and statistical power when true parameters are zero or nonzero, respectively. **(B)** Type I error when the true genetic correlation is 0. **(C)** qq-plot of p value when the true genetic correlation is set to be 0.

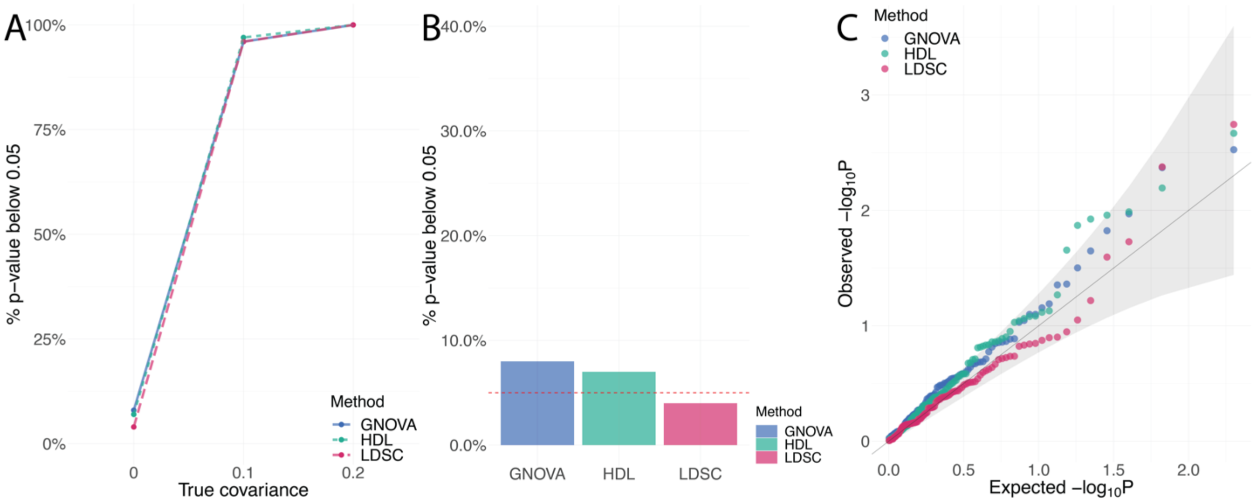

**Supplementary Figure 6. Comparisons of genetic covariance and correlation estimation using external reference panel with matched LD under model misspecification.** We set 10% of the total SNPs as the causal SNPs to simulate the phenotypes. The estimates for **(A)** genetic covariance and **(B)** genetic correlation among LDSC, GNOVA, and HDL are summarized by boxplots which show the quantiles of the estimates. The red dashed lines represent true values.

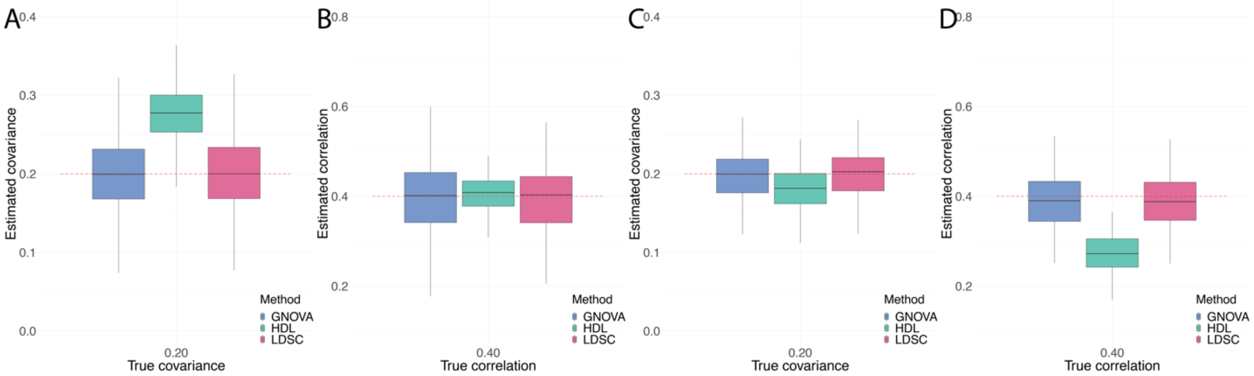

**Supplementary Figure 7. Addition simulations for HDL that used 99.99% SNPs in the reference panel to simulate phenotypes.** We used external refence panel with matched LD. Panels A-B show the estimation of HDL for **(A)** genetic covariance and **(B)** genetic correlation when the two GWASs were simulated on the same dataset with a 100% sample overlap. Panels C-D show the estimation of HDL for **(C)** genetic covariance and **(D)** genetic correlation when the GWASs were simulated on two non-overlapping datasets. The red dashed lines represent true values.

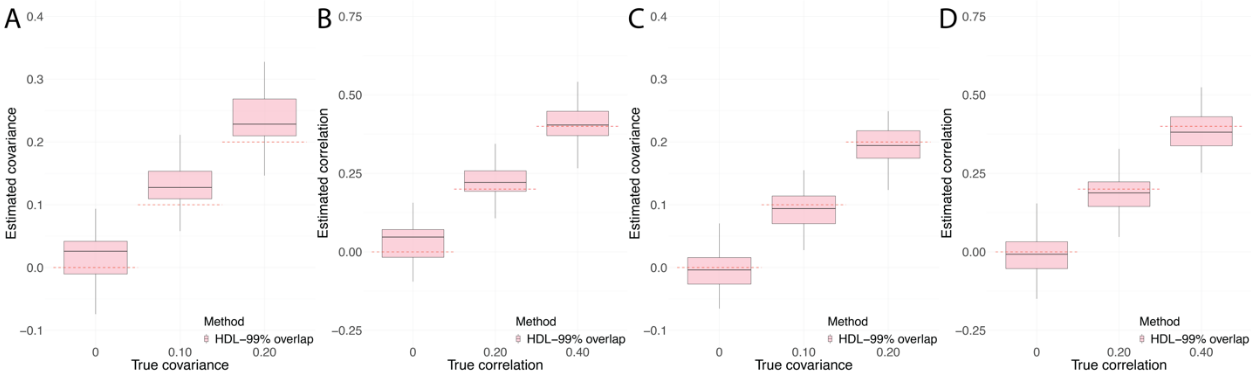

**Supplementary Figure 8. Comparisons of genetic covariance and correlation estimation using external reference panel with mismatched LD under model misspecification.** We set 10% of the total SNPs as the causal SNPs to simulate the phenotypes. The estimates for **(A)** genetic covariance and **(B)** genetic correlation among LDSC, GNOVA, HDL, and REML are summarized by boxplots which show the quantiles of the estimates. The red dashed lines represent true values.

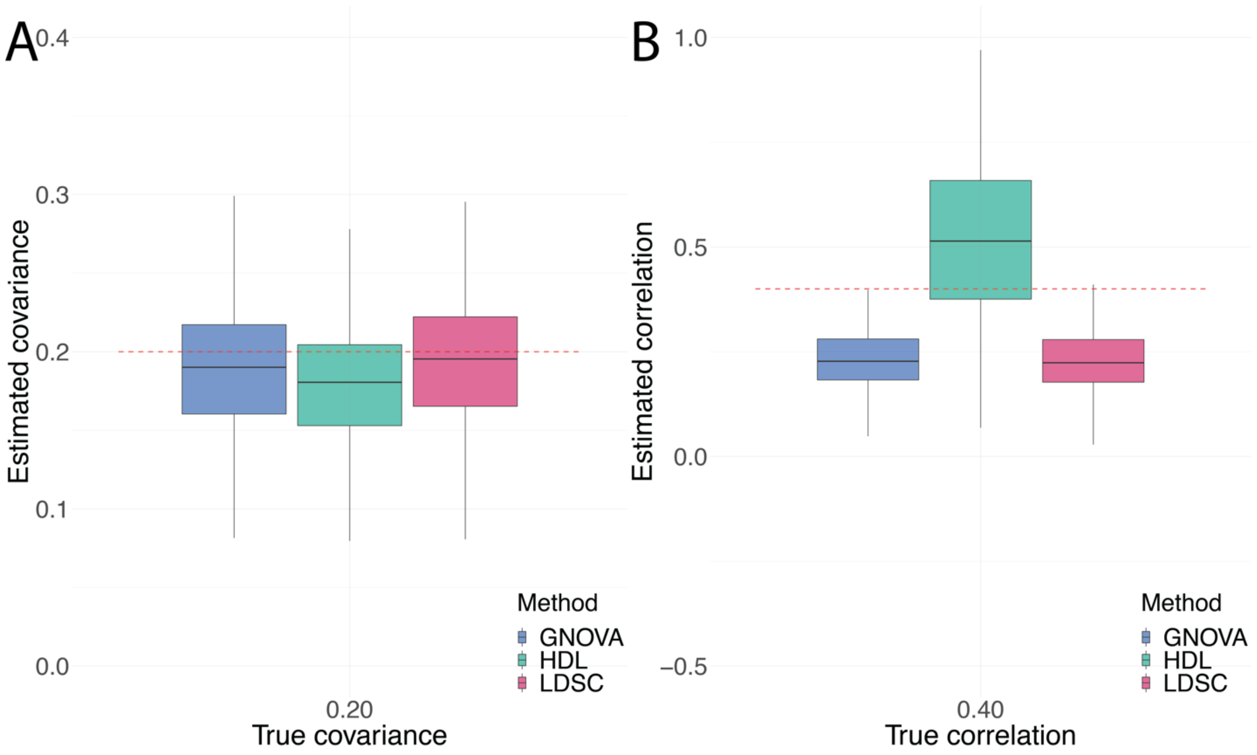

**Supplementary Figure 9. Evaluation of type I error control and statistical power among genetic correlation (covariance) estimation methods using external reference panel with mismatched LD.** **(A)** Type I error and statistical power when true parameters are zero or nonzero, respectively. **(B)** Type I error when the true genetic correlation is 0. **(C)** qq-plot of p value when the true genetic correlation is set to be 0.

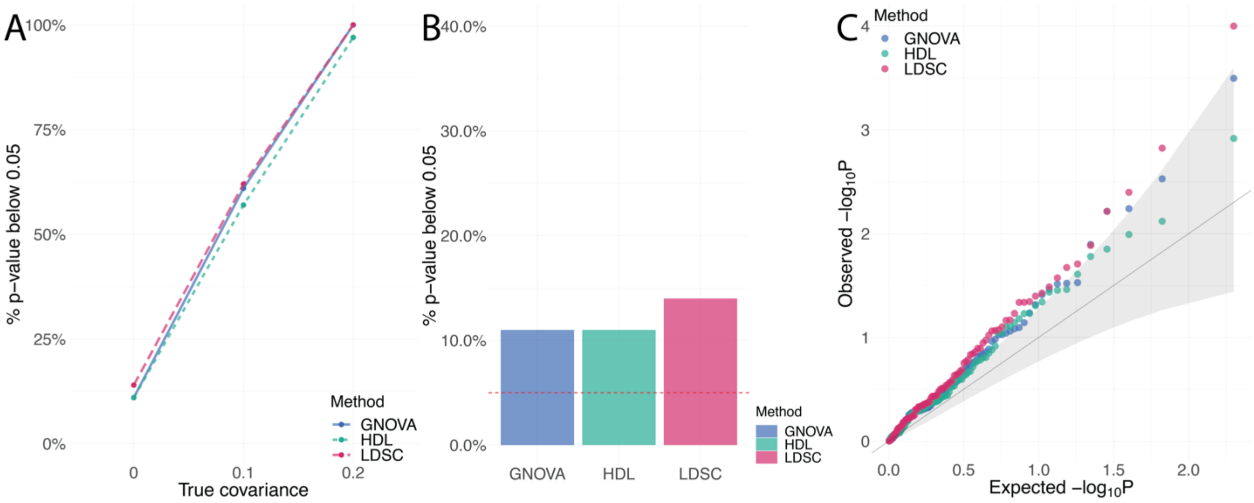

**Supplementary Figure 10. Comparison of genetic correlation estimation on binary traits using in-sample reference panel.** The threshold in the liability model to simulate binary traits was set to be $\Phi^{-1}(0.8) = 0.84$. The estimated genetic correlations by LDSC, GNOVA, HDL, and REML are summarized by boxplots which show the quantiles of the estimates. The red dashed lines represent true values.

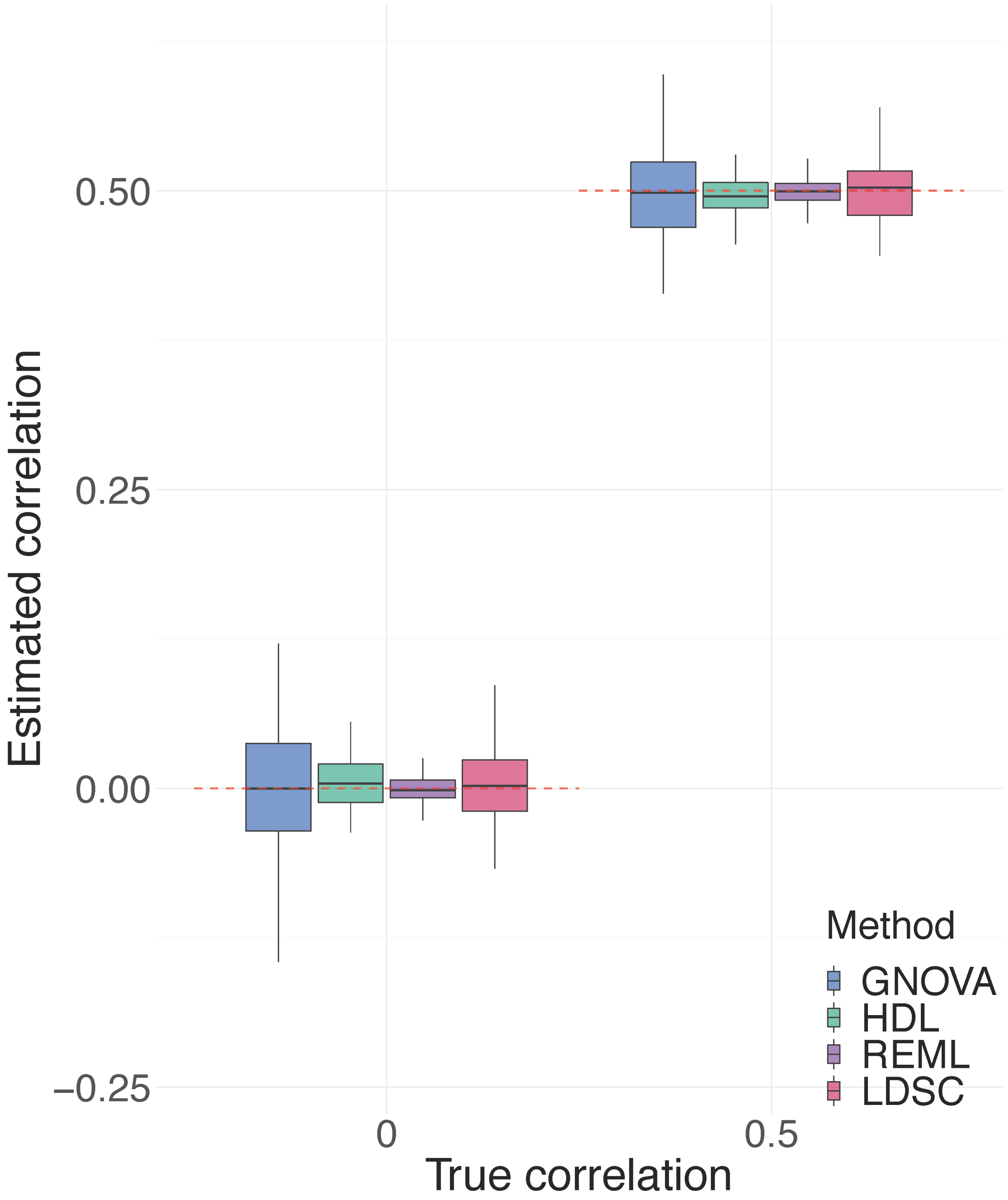

**Supplementary Figure 11. Comparisons of genetic correlation estimation on binary traits using external reference panel with matched LD.** The threshold in the liability model to simulate binary traits was set to be $\Phi^{-1}(0.8) = 0.84$. Genetic correlation among LDSC, GNOVA, HDL and REML are demonstrated by boxplot which shows the quantiles of the estimates. The red dashed lines represent true values. Panel A-B compare the estimation of genetic correlation when the two GWASs were simulated on the same dataset with **(A)** a 100% sample overlap and **(B)** two non-overlapping datasets.

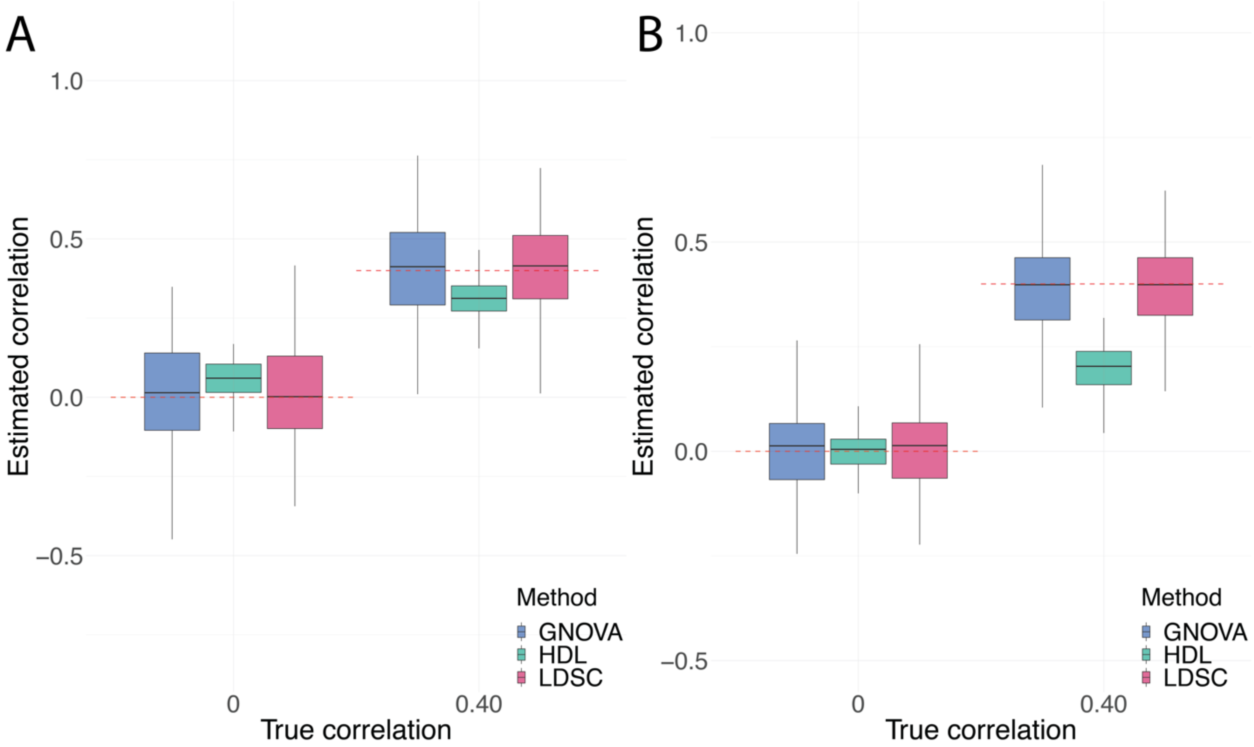

**Supplementary Figure 12. Comparison of genetic correlation estimation on binary traits using external reference panel with mismatched LD.** The threshold in the liability model to simulate binary traits was set to be $\Phi^{-1}(0.8) = 0.84$. Genetic correlation among LDSC, GNOVA, HDL and REML are demonstrated by boxplot which shows the quantiles of the estimates. The red dashed lines represent true values.

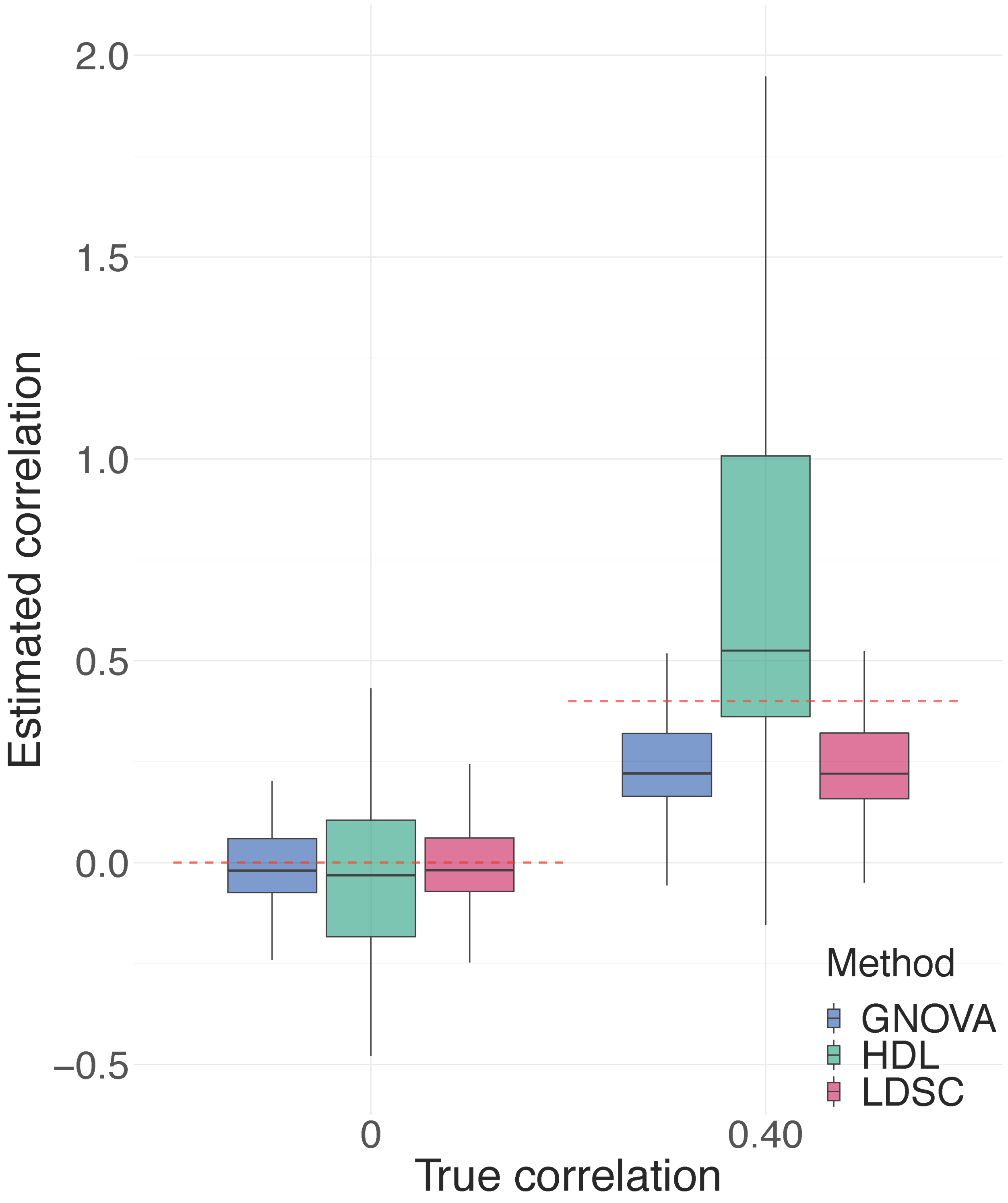

**Supplementary Figure 13. Evaluation of type I error control and statistical power among genetic correlation (covariance) estimation methods using in-sample reference panel and applying to binary traits.** The threshold in the liability model to simulate binary traits was set to be $\Phi^{-1}(0.8) = 0.84$. **(A)** Type I error and statistical power when true parameters are zero or nonzero, respectively. **(B)** Type I error when the true genetic correlation is 0. **(C)** qq-plot of p value when the true genetic correlation is set to be 0.

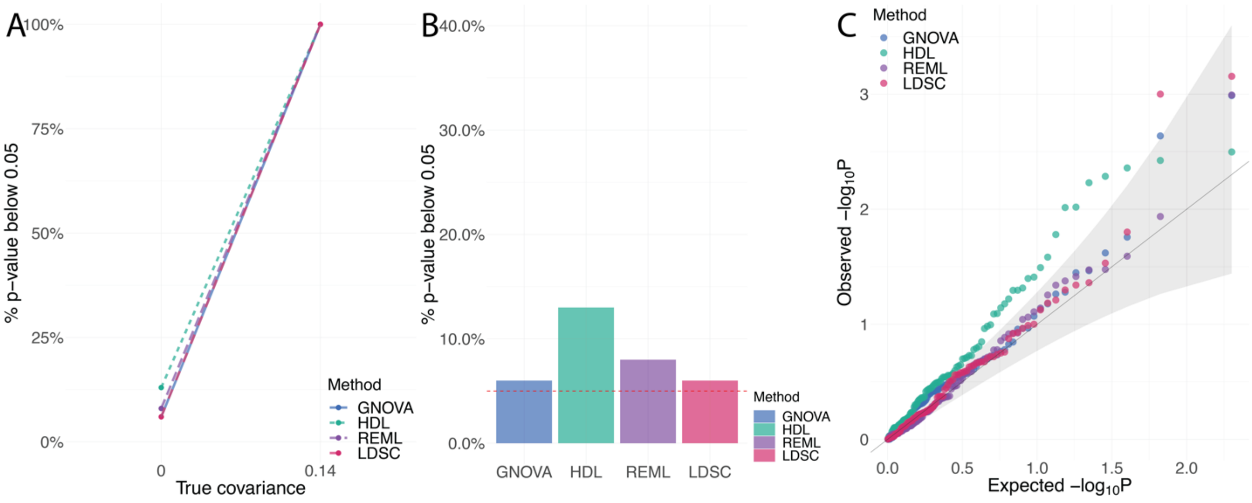

**Supplementary Figure 14. Evaluation of type I error control and statistical power among genetic correlation (covariance) estimation methods using external reference panel and applying to binary traits with matched LD on completely overlapping dataset (set 1).** The threshold in the liability model to simulate binary traits was set to be $\Phi^{-1}(0.8) = 0.84$. **(A)** Type I error and statistical power when true parameters are zero or nonzero, respectively. **(B)** Type I error when the true genetic correlation is 0. **(C)** qq-plot of p value when the true genetic correlation is set to be 0.

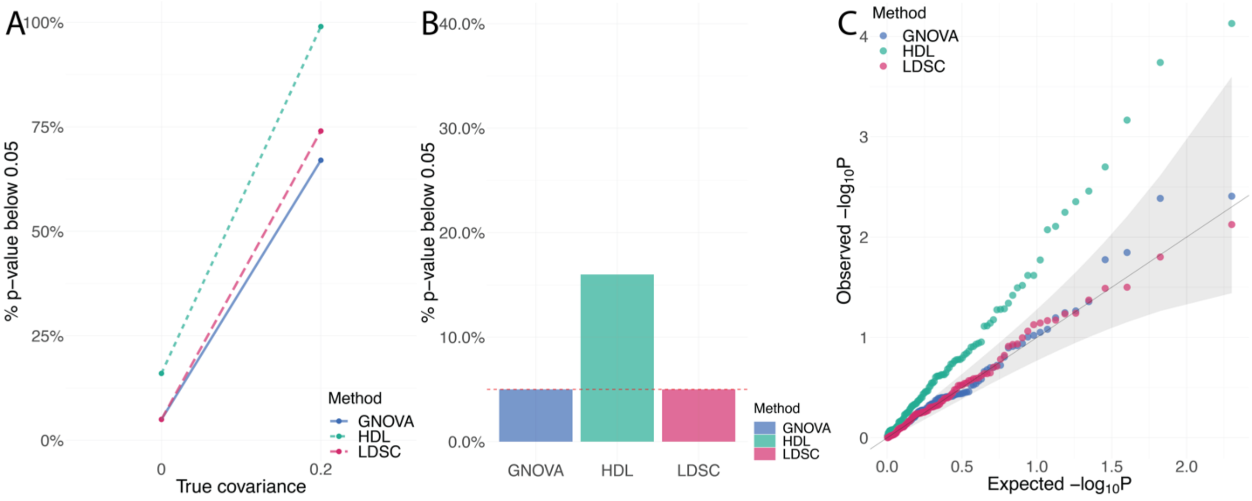

**Supplementary Figure 15. Evaluation of type I error control and statistical power among genetic correlation (covariance) estimation methods using external reference panel and applying to binary traits with matched LD on non-overlapping dataset (set 1 and set 2).** The threshold in the liability model to simulate binary traits was set to be $\Phi^{-1}(0.8) = 0.84$. **(A)** Type I error and statistical power when true parameters are zero or nonzero, respectively. **(B)** Type I error when the true genetic correlation is 0. **(C)** qq-plot of p value when the true genetic correlation is set to be 0.

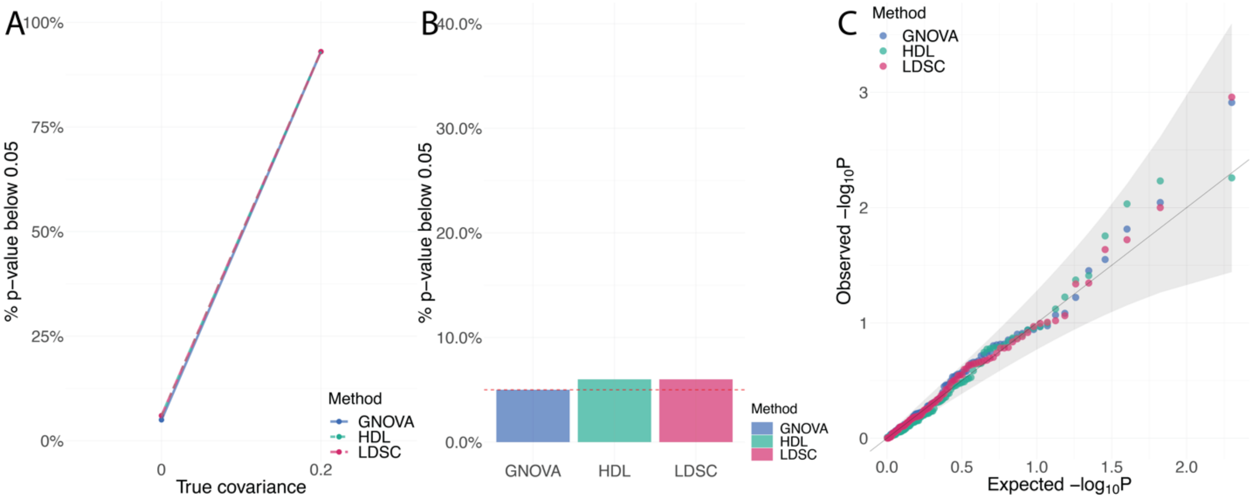

**Supplementary Figure 16. Evaluation of type I error control and statistical power among genetic correlation (covariance) estimation methods using external reference panel and applying to binary traits with mismatched LD.** The threshold in the liability model to simulate binary traits was set to be $\Phi^{-1}(0.8) = 0.84$. **(A)** Type I error and statistical power when true parameters are zero or nonzero, respectively. **(B)** Type I error when the true genetic correlation is 0. **(C)** qq-plot of p value when the true genetic correlation is set to be 0.

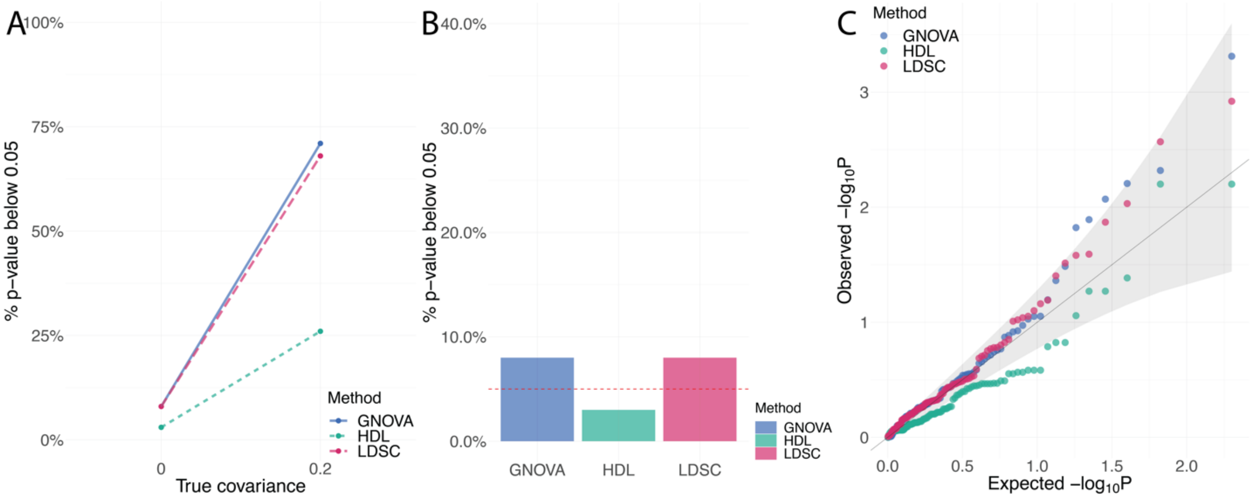

**Supplementary Figure 17. Comparisons of genetic covariance and correlation estimation between REML and summary-data-based methods on UKBB traits.** The estimates of genetic **(A)** covariance and **(B)** correlation are summarized by scatter plots. The R squares of LDSC, GNOVA and HDL for genetic covariance are 0.99, 0.96, and 0.99, respectively. The R squares of LDSC, GNOVA and HDL for genetic correlation are 0.99, 0.98, and 0.99, respectively. The color and shape of each point represent the method. The dashed lines are $y = x$. Genetic correlation estimation showed more consistency between REML and summary-data-based methods than genetic covariance estimation.

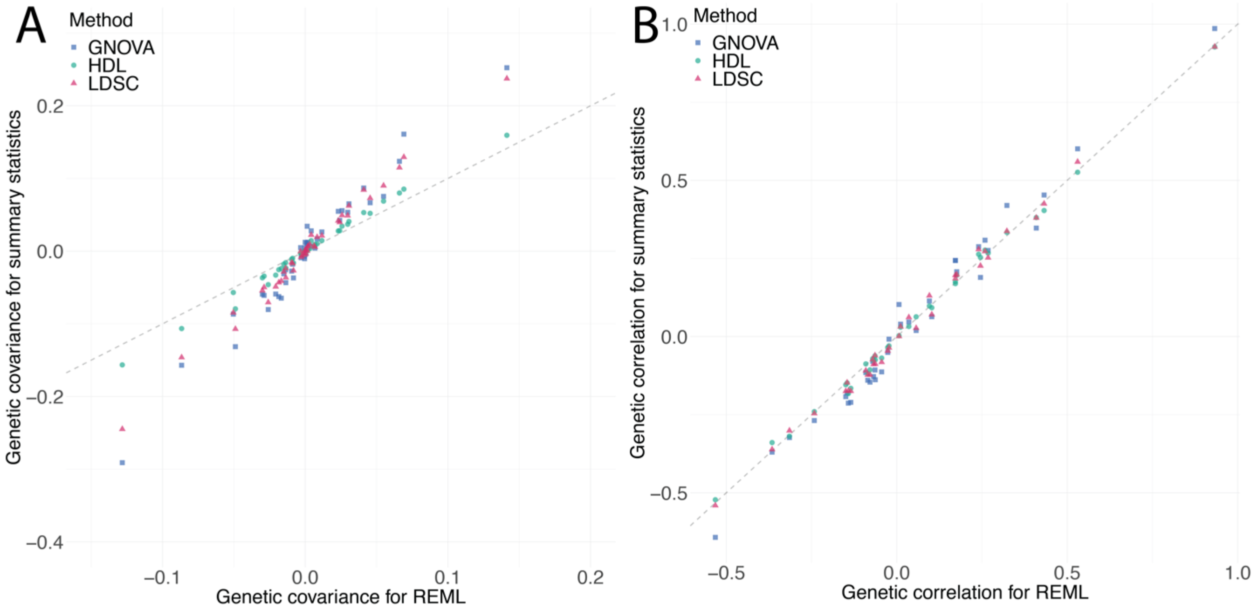

**Supplementary Figure 18. Comparisons of genetic covariance and correlation estimation between REML and summary-data-based methods on WTCCC and NFBC traits.** The estimates of genetic **(A)** covariance and **(B)** correlation are summarized by scatter plots. The R squares of LDSC, GNOVA and HDL for genetic covariance are 0.58, 0.47, and 0.52, respectively. The R squares of LDSC, GNOVA and HDL for genetic correlation are 0.66, 0.54, and 0.50, respectively. The color and shape of each point represent the method. The dashed lines are $y = x$. Genetic correlation estimation showed more consistency between REML and summary-data-based methods than genetic covariance estimation.

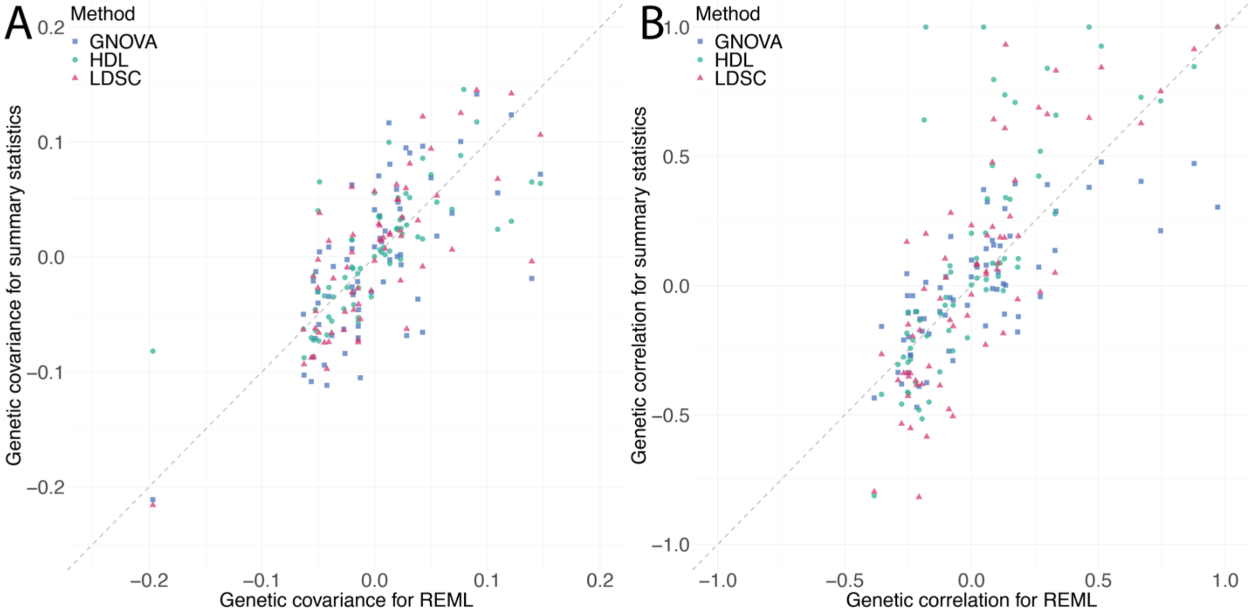

**Supplementary Figure 19. Genetic correlation among 30 complex traits.** The results of **(A)** GNOVA, **(B)** HDL, and **(C)** LDSC are presented by heatmaps. Asterisks highlight significant genetic correlations after Bonferroni correction for 435 pairs. We grouped traits with hierarchical clustering applied to genetic correlations. We summarized detailed information about each trait, including abbreviations, in **Supplementary Table 1**.

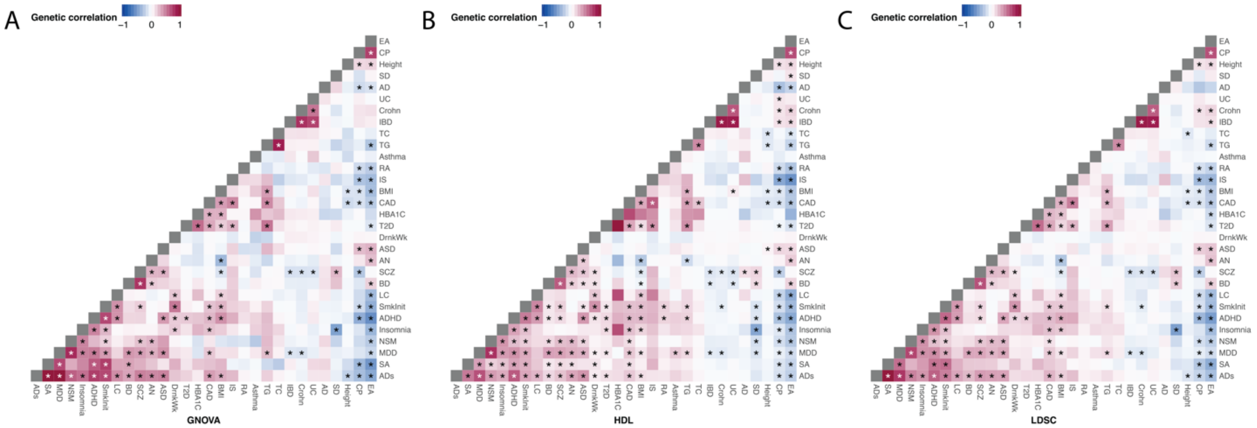

**Supplementary Figure 20. Trait pairs with significant genetic correlation identified by LDSC, GNOVA, and HDL ($p < 1.15e-4$).** This plot uses bars to break down the Venn diagram of overlapped regions in different categories. The three categories shown in the lower panel are correlated trait pairs identified by LDSC, GNOVA and HDL.

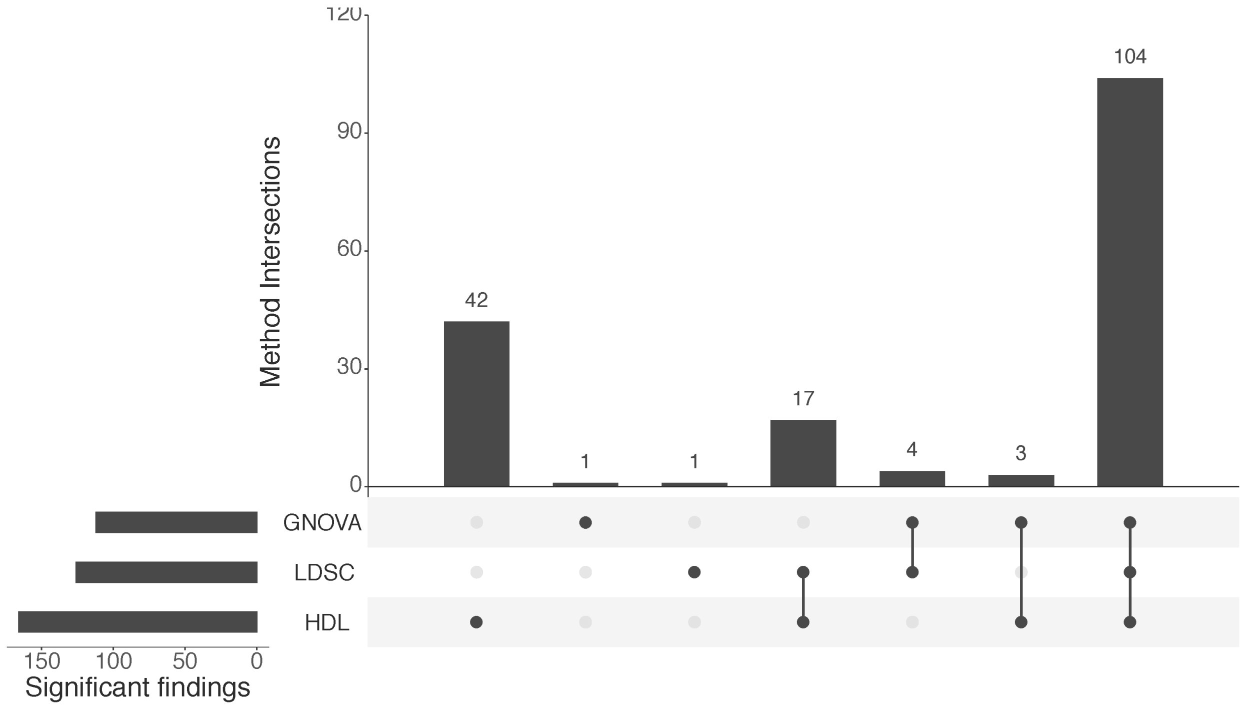

**Supplementary Figure 21. Volcano plot for genetic correlation.** The relation between -log10(Pvalue) and genetic correlation estimation for **(A)** GNOVA, **(B)** HDL and **(C)** LDSC are shown. Each point represents a trait pair. Color of each data point represents the significance and direction of global correlation.

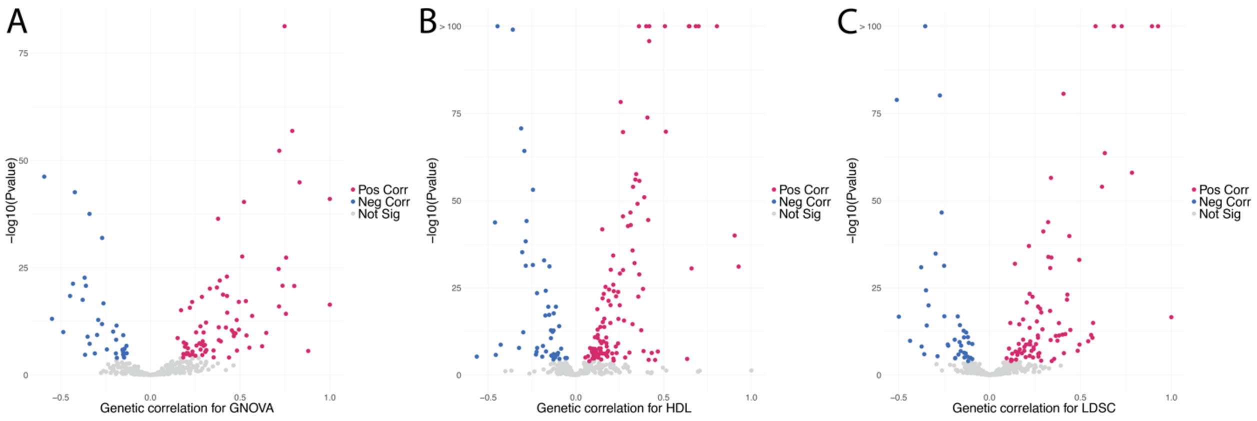
